# ISL1 is necessary for auditory neuron development and contributes towards tonotopic organization

**DOI:** 10.1101/2021.09.03.458707

**Authors:** Iva Filova, Kateryna Pysanenko, Mitra Tavakoli, Simona Vochyanova, Martina Dvorakova, Romana Bohuslavova, Ondrej Smolik, Valeria Fabriciova, Petra Hrabalova, Sarka Benesova, Lukas Valihrach, Jiri Cerny, Ebenezer N. Yamoah, Josef Syka, Bernd Fritzsch, Gabriela Pavlinkova

**Author notes:** These authors contributed equally: Iva Filova, Kateryna Pysanenko.

## Abstract

A cardinal feature of the auditory pathway is frequency selectivity, represented in a tonotopic map from the cochlea to the cortex. The molecular determinants of the auditory frequency map are unknown. Here, we discovered that the transcription factor ISL1 regulates the molecular and cellular features of auditory neurons, including the formation of the spiral ganglion and peripheral and central processes that shape the tonotopic representation of the auditory map. We selectively knocked out *Isl1* in auditory neurons using *Neurod1^Cre^* strategies. In the absence of *Isl1*, spiral ganglion neurons migrate into the central cochlea and beyond, and the cochlear wiring is profoundly reduced and disrupted. The central axons of *Isl1* mutants lose their topographic projections and segregation at the cochlear nucleus. Transcriptome analysis of spiral ganglion neurons shows that *Isl1* regulates neurogenesis, axonogenesis, migration, neurotransmission-related machinery, and synaptic communication patterns. We show that peripheral disorganization in the cochlea affects the physiological properties of hearing in the midbrain and auditory behavior. Surprisingly, auditory processing features are preserved despite the significant hearing impairment, revealing central auditory pathway resilience and plasticity in *Isl1* mutant mice. Mutant mice have a reduced acoustic startle reflex, altered prepulse inhibition, and characteristics of compensatory neural hyperactivity centrally. Our findings show that ISL1 is one of the obligatory factors required to sculpt auditory structural and functional tonotopic maps. Still, upon *Isl1* deletion, the ensuing central compensatory plasticity of the auditory pathway does not suffice to overcome developmentally induced peripheral dysfunction of the cochlea.

## Introduction

Spiral ganglion neurons (SGNs) are bipolar, extending peripheral processes to the hair cells within the sensory epithelium (the organ of Corti) and central axons towards the cochlear nucleus (CN) complex, the first auditory nuclei in the brain. Sound-induced vibrations that reach the cochlea are amplified by the outer hair cells (OHCs) organized in three rows and innervated by type II SGNs. The inner hair cells (IHCs) receive, transduce, and transmit the auditory signal to type I SGNs that convey the signal via the vestibulocochlear cranial nerve to the CN of the brainstem. The auditory neurons are organized within the cochlea in an orderly fashion according to frequency, with high frequencies at the base and low frequencies at the apex ^1, 2^. The tonotopic organization of type I SGNs corresponds to multiple diversities in their molecular profiles, connectivity patterns, and physiological features along the tonotopic axis ^3, 4, 5^. The cochleotopic or tonotopic pattern is maintained throughout the auditory pathways in the brain ^6^. The central auditory pathway transmits ascending acoustic information from the CN through the lateral lemniscus complex, the inferior colliculus in the midbrain, the medial geniculate nucleus of the thalamus to the auditory cortex ^7^. The efferent motor neurons consist of the medial olivocochlear motor neurons, which modulate cochlear sound amplification by OHCs. In contrast, lateral olivocochlear motor neurons innervate afferent type I sensory neurons and regulate cochlear nerve excitability ^7, 8, 9^.

The cellular and molecular regulation of neuronal migration and the establishment of tonotopic connections to the hair cells or neurons of the hindbrain’s first auditory nuclei, the CN, are not fully understood. Several transcription factors govern the development of inner ear neurons, including NEUROG1 ^10^, NEUROD1 ^11^, GATA3 ^12, 13^, and POU4F1 ^14^. The transcription factor NEUROD1 is vital for the differentiation and survival of inner ear neurons ^15, 16, 17^. We previously demonstrated that *Isl1^Cre^*-mediated *Neurod1* deletion (*Neurod1CKO*) results in a disorganized cochleotopic projection from SGNs ^18^, affecting acoustic information processing in the central auditory system of adult mice at the physiological and behavioral levels ^19^. During inner ear development, ISL1 is expressed in the differentiating neurons and sensory precursors ^18, 20, 21, 22, 23^. *Isl1* is expressed in all four populations of SGNs (type Ia, Ib, Ic, and type II) identified by single-cell RNA sequencing ^4^. Studies suggest that ISL1 plays a role in developing neurons and sensory cells, but there has been no direct evaluation of ISL1 function in the inner ear. Using *Neurod1^Cre^* ^24^, we conditionally deleted *Isl1* (*Isl1CKO*) in neurons without directly affecting the development of the inner ear sensory epithelium.

This work provides the first genetic and functional evidence for ISL1’s role in establishing the spiral ganglion peripheral projection map and proper central auditory circuitry. Most *Isl1CKO* neurons migrated into the center of the cochlear modiolus. They extended outside the cartilaginous/bony otic capsule, in contrast, to control animals with an arrangement of SGNs in parallel to the spiraling cochlear duct. Additionally, we analyzed the transcriptome of neurons, hearing function, sound information processing in the inferior colliculus, and auditory behavior of *Isl1CKO* to demonstrate how abnormal SGN development affects the formation, wiring, and operation of the auditory sensory maps.

## Results

### ISL1 controls neuronal phenotype in the cochlea

To investigate the role of ISL1 in inner ear neuron development, we eliminated *Isl1* by crossing *Isl1^loxP/loxP^* mice ^25^ with *Neurod1^Cre^* mice ^24^. *Neurod1^Cre^* is expressed in sensory neurons but not in the sensory epithelium, as visualized by *tdTomato* reporter expression (Supplementary Fig. 1). Analyses of ISL1 expression in *Isl1CKO* confirmed efficient recombination by *Neurod1^Cre^* with virtually no expression of ISL1 during the differentiation of neurons in the inner ear ganglion as early as embryonic day E10.5, and later in the cochlear neurons, shown at E14.5 and postnatal day 0 (P0), in contrast to high levels of ISL1 in neurons in the control inner ear (Supplementary Fig. 2). It is worth mentioning that no difference was observed in the density of Schwann cells (Supplementary Fig. 2e, f).

The organization of the sensory epithelium in the organ of Corti of *Isl1CKO* was comparable to controls with three rows of OHCs and one row of IHCs (Fig. 1a-f), as shown by immunolabeling with the hair cell differentiation marker, myosin VIIA (Myo7a). Whole-mount anti-tubulin staining of innervation showed reduced and disorganized radial fibers. Significant gaps were found between radial fiber bundles, crisscrossing fibers, and an unusual dense innervation in the apex, with some large fiber bundles bypassing the organ of Corti and extending to the lateral wall in the *Isl1CKO* cochlea (arrowhead, Fig. 1f).

**Figure 1.**
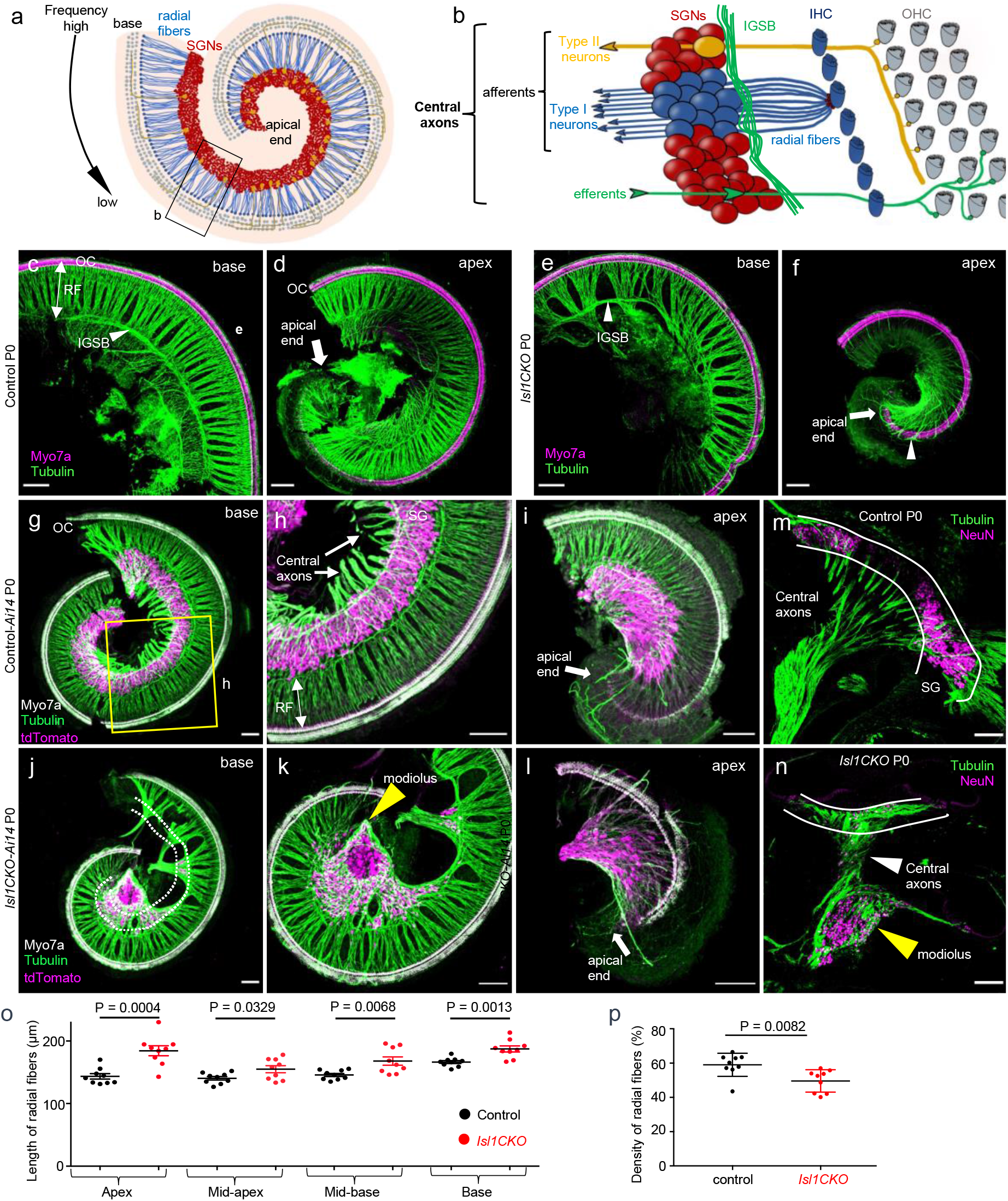
*Isl1* deletion results in abnormalities in cochlear innervation and the formation of the spiral ganglion. (**a**) Diagram of the organization of the cochlea. Spiral ganglion neurons (SGNs) are organized tonotopically from the apex to the base of the cochlea. High-frequency sounds maximally stimulate the base of the cochlea, whereas the largest response to low-frequency sounds occurs in the cochlear apex. **(b)** The top view diagram onto the sensory epithelium shows outer hair cells (OHCs), inner hair cells (IHCs), and SGNs. Type I neurons extend radial fibers (RF) toward IHCs (5-30 neurons innervate one IHC), and type II neurons (representing 5% of SGNs) receive input from OHCs. The central processes of SGNs relay acoustic information to the brain. Efferent axons from the superior olivary complex, forming the intraganglionic spiral bundle (IGSB), innervate OHCs. **(c-f)** Representative images of whole-mount immunolabeling of the base and apex with anti-Myo7a (hair cell marker) and anti-α-tubulin (nerve fibers) show reduced and disorganized RF and missing or altered efferent fibers forming IGSB (arrowhead) in *Isl1CKO* compared to control. Abnormalities in innervation, including larger gaps between radial fiber bundles, and fiber bundles bypassing OC (arrowhead in f), are noticeable in *Isl1CKO*. **(g-i)** The shape of the spiral ganglion (SG) is shown in the basal and apical half of the whole-mounted cochlea of control-*Ai14* reporter mice at P0. TdTomato^+^ neurons form a spiral located in the Rosenthal’s canal and parallel to the OC (anti-Myo7a labeled hair cells), nerve fibers are labeled with anti-tubulin. **(j-l)** In *Isl1CKO*, only some tdTomato^+^ neurons are in the Rosenthal’s canal (delineated by white dotted lines), but many neurons are in the centre of the cochlea, a conical-shaped structure, the modiolus (yellow arrowhead). **(m, n)** The vibratome sections of the cochlea labeled by anti-NeuN (a nuclear marker of differentiated neurons) and anti-tubulin (nerve fibers) show the unusual position of cochlear neurons in the modiolus, and neurons entwined in central axons in *Isl1CKO* compared to control (white lines delineate a normal position of the SG). **(o)** The length of radial fibers was measured in the confocal images from cochlear whole-mount preparations in the apex, mid-apex, mid-base, and base. Data are expressed as mean ± SEM (n = 3 radial fiber bundles/each region/3 cochlea/genotype); unpaired *t*-test. **(p)** Fiber density was quantified in the base, mid-base, and mid-apex (n = 3 mice per genotype). Error bars represent mean ± SD; unpaired *t*-test. Scale bars: 100 μm.

Next, we evaluated the effects of *Isl1* deletion on the formation of the spiral ganglion. The SGN somas were well restricted to the Rosenthal’s canal in control animals, arranged in a spiral parallel to the cochlear duct (Fig. 1g-i). In contrast, unevenly distributed neurons were found in *Isl1CKO* with many SGNs accumulated beyond the Rosenthal’s canal, in the conical central part of the cochlea, the modiolus (Fig. 1j-l), which usually carries only afferent and efferent fibers. The unusual accumulation of cochlear neurons intertwined with the central axons in the modiolus of *Isl1CKO* compared to the control mice is depicted in the immunolabeled sections of the cochlea (Fig. 1m, n). To determine whether neuron number was affected by apoptosis during inner ear development, we performed immunostaining for cleaved Caspase-3 (Supplementary Fig. 3). We found moderately increased apoptosis of neurons in *Isl1CKO* at E12.5 but not at E10.5, although the size of inner ear ganglia was unaffected at E12.5. These results imply that missing neurons in the Rosenthal’s canal are due to the altered migration of SGNs rather than neuronal death in the developing inner ear. Correspondingly, most apoptotic neurons were found outside the cochlea in the vestibular ganglion area in E14.5 *Isl1CKO*. In line with the unusual position of SGNs in the *Isl1CKO* cochlea, the radial fibers were significantly lengthened in all regions of the *Isl1CKO* cochlea (Fig. 1o), but the overall radial fiber density was reduced (Fig. 1p).

Interestingly, *Isl1* deficiency caused a shortening of the cochlea, which was on average ~20% and 25% shorter at P0 and in adults, respectively (Fig. 2a, b). Since *Neurod1^Cre^* is expressed only in neurons, the shortening of the cochlea appears to be a secondary effect of the abnormalities in neuronal development. A similar phenotype of truncated growth of the cochlea was reported for *Neurod1* and *Neurog1* mutants because of alternations in spatiotemporal gene expression ^10, 18, 19, 26^. Correspondingly to the truncated cochlear phenotype ^10, 18, 19, 26^, detailed analyses showed abnormalities in the epithelium at the apical end with disorganized rows of OHCs and ectopic IHCs among OHCs in the *Isl1CKO* cochlea (Fig. 2c). The disorganized apical epithelium represented ~11.8% of the length of the organ of Corti in adults (n = 6). Anti-tubulin labeling revealed a reduced and disorganized innervation in the region of OHCs in *Isl1CKO* (Fig. 2d). In the controls, fibers extending through the tunnel of Corti were oriented towards the base to form three parallel outer spiral bundles. In contrast, guiding defects in the extension of these fibers in *Isl1CKO* were apparent, with neurites extending with random turns towards the base and the apex (arrowheads in Fig. 2d). To further investigate cochlear histopathology in *Isl1CKO*, we evaluated synaptic contacts from type I SGNs onto IHCs in 2-month-old mice (Fig. 2e). CtBP2^+^ ribbon synapse number was reduced in *Isl1CKO,* representing ~46%, 51%, 56%, and 69% ribbons of the control base, mid-base, mid-apex, and apex, respectively (Fig. 2f, Supplementary Table 1).

**Figure 2.**
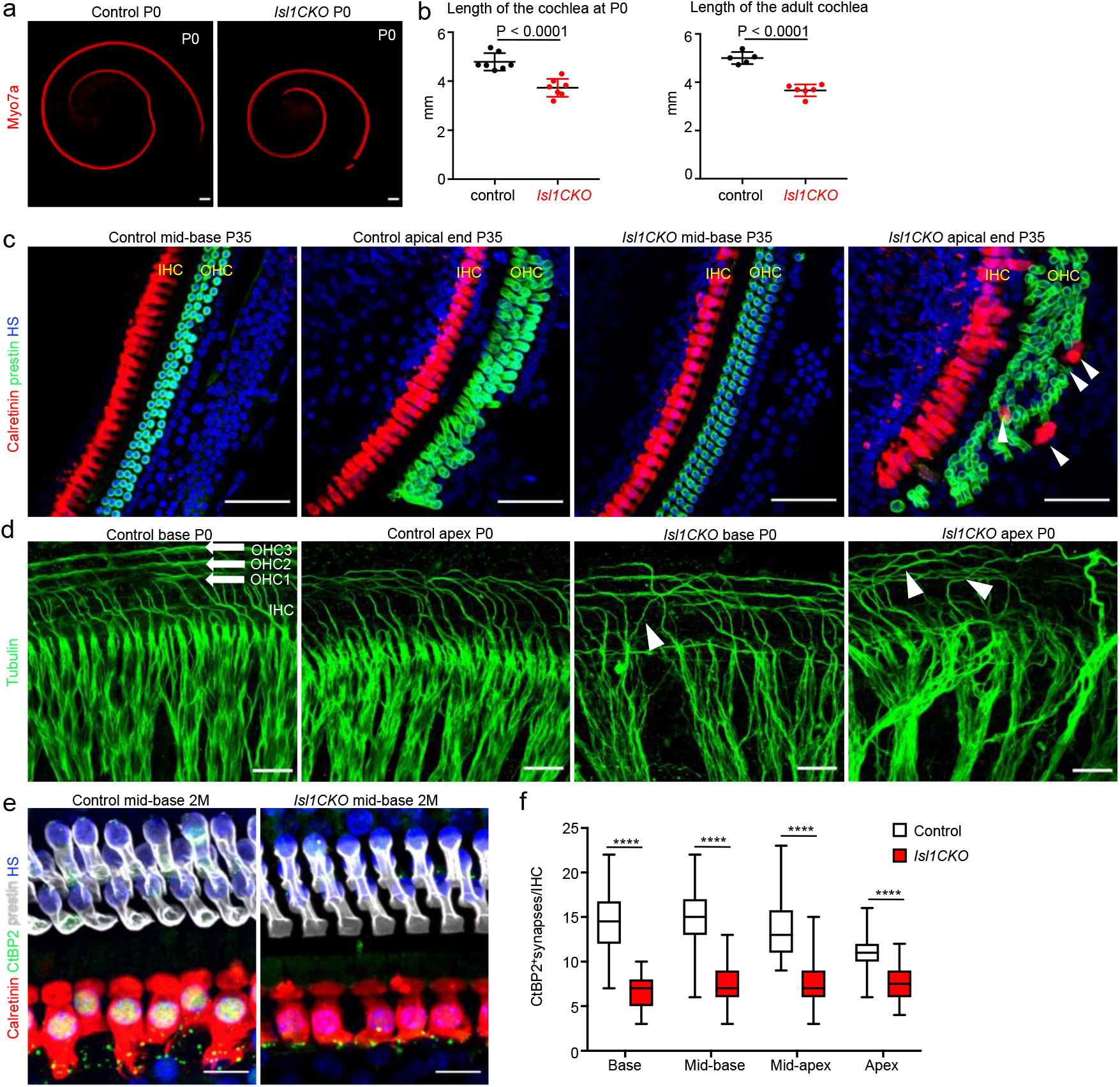
The length of the organ of Corti is shortened, the apical end sensory epithelium is disorganized, and presynaptic ribbons are reduced in *Isl1CKO*. (**a**) The organ of Corti of *Isl1CKO* is shorter than control, as shown by anti-Myo7a labeling of hair cells in the cochlea at P0. **(b)** The length of the organ of Corti was measured at postnatal day P0 and in 2-month-old mice. Error bars represent mean ± SD; unpaired *t*-test. (**c**) Immunohistochemistry for prestin (a marker for OHCs) and calretinin (a marker for IHCs) shows a comparable cytoarchitecture of the organ of Corti in the cochlear mid-base of controls and *Isl1CKO* but disorganized rows of OHCs in the apical end with multiple OHC rows and ectopic calretinin^+^ IHCs among OHCs (arrowheads) in *Isl1CKO* at P35. **(d)** Higher-magnification images show a detail of anti-tubulin labeled innervation with fibers forming three parallel outer spiral bundles (arrows) that innervate multiple OHCs and turn toward the base in the control cochlea. Guiding defects in the extension of these fibers to OHCs are obvious in *Isl1CKO* with some fibers randomly turned toward the apex (arrowheads); note the disorganization of radial fiber bundles in *Isl1CKO*. **(e)** Representative images of whole-mount immunolabeling of the cochlear mid-base with calretinin (IHCs) and anti-CtBP2 (presynaptic ribbons) of 2-month-old mice. **(f)** CtBP2^+^ synapses were counted in 10 IHCs per each cochlear region (n = 6 mice per genotype). Box plots indicate median (middle line), 25th, 75th percentile (box) and min to max (whiskers); multiple *t*-tests, ****P < 0.000001. HS, Hoechst nuclear staining; IHC, inner hair cell; OHC, outer hair cell. Scale bars: 100 μm (a), 50 μm (c), 20 μm (d), 10 μm (e).

Altogether, these results demonstrate the prominent effects of *Isl1* deletion on the formation of the spiral ganglion and innervation patterns in the cochlea. The truncated cochlea in *Isl1CKO* indicates an altered spatial base-to-apex organization of structural and physiological characteristics of auditory neurons.

### ISL1 regulates neuronal identity and differentiation programs of SGNs

To gain insight at the molecular level on how the *Isl1* elimination causes the neuronal phenotype in the cochlea, we sought to identify potential ISL1 targets through global transcriptome analysis. We opted to use Bulk-RNA sequencing to obtain sequencing depth and high-quality data ^27^. Six biological replicates were used per genotype, and each replicate contained a total of 100 tdTomato^+^ SGNs isolated from the E14.5 cochlea. Spiral ganglia were dissected, dissociated into single cells, and fluorescent tdTomato^+^ cells were FACS-sorted (Fig. 3a). Compared to controls, 650 protein-coding genes were differentially expressed in *Isl1CKO* neurons (adjusted p-value < 0.05, and fold change > 2 cut-off values, see Methods), 332 genes down- and 318 genes up-regulated (Fig. 3b, Supplementary Data 1). Gene ontology (GO) term enrichment analysis for the GO term category biological process revealed highly enriched GO terms associated with neuron development, including “neurogenesis”, “neuron differentiation”, and “nervous system development”, in both up- and down-differentially expressed genes (Fig. 3c, Supplementary Data 2). The most enriched and specific GO categories for downregulated genes were associated with neurotransmission-related machineries, such as “transmembrane transporter”, “voltage-gated channel”, “cation and ion transport”, and “membrane potential regulation”, indicating changes in neuronal cell functions. The analysis identified enrichment of downregulated genes involved in axon development, guidance, axonogenesis, and neuronal migration. These genes included members of all four classes of axon guidance molecules and their cognate receptors, the ephrins-Eph, semaphorins-plexin, netrin-unc5, and slit-roundabout ^28^, for instance, *Epha5, Epha4, Pdzrn3, Sema3e, Sema6d, Plxna4, Ntn3, Ntng1, Ntng2, Kirrel3, Unc5b, Unc5c, Slitrk3*, and *Slitrk1*. The neurotrophic tyrosine kinase receptors (*Ntrk2* and *Ntrk3*) and a G-protein-coupled chemokine receptor (*Cxcr4*), important regulators of neuronal migration, were also downregulated ^29^.

**Figure 3.**
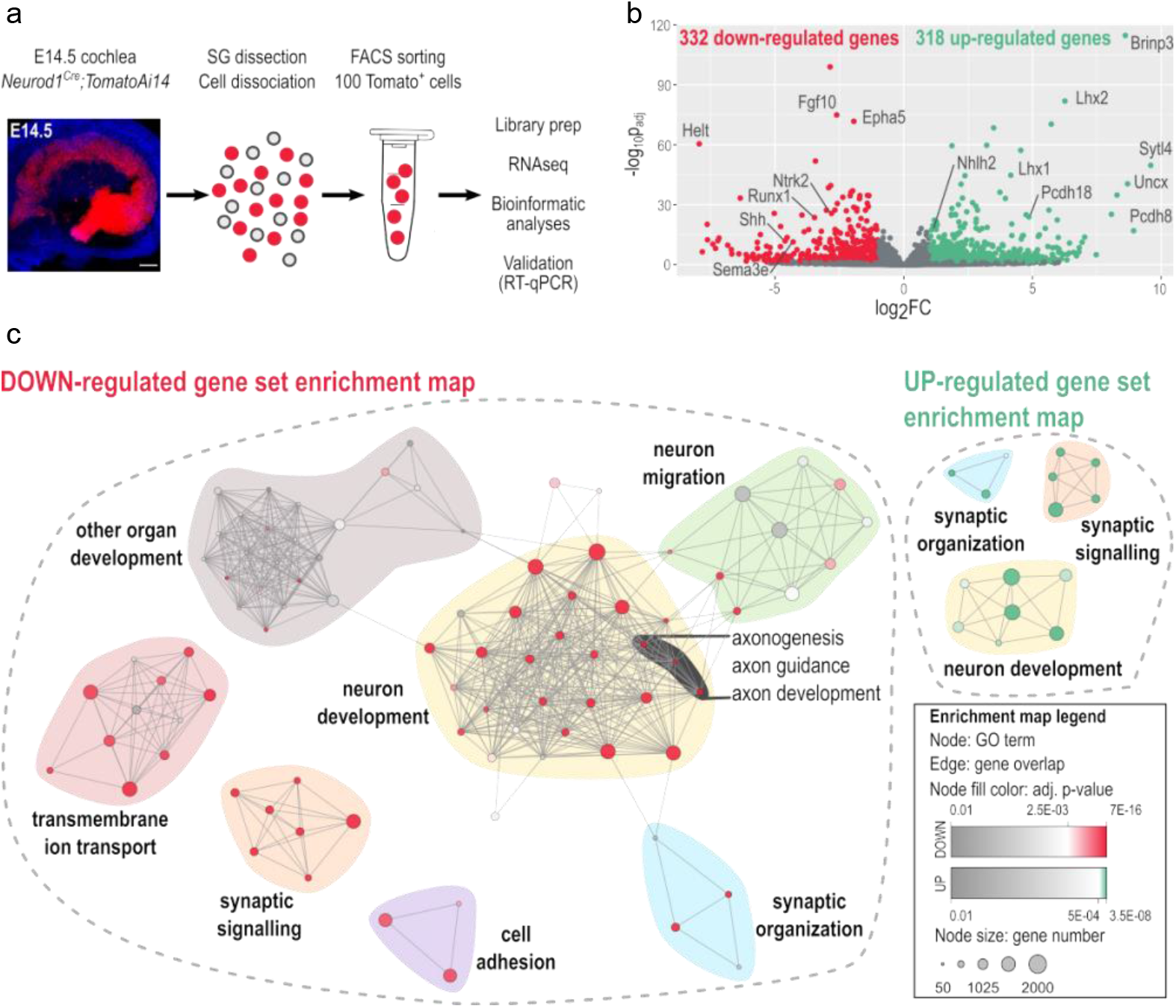
ISL1-mediated transcription signature in cochlear neurons. **(a)** Image of the whole-mount cochlea with genetically labeled tdTomato neurons in the spiral ganglion at E14.5 [Hoechst staining (blue), Scale bar: 100 μm]; and workflow depicts microdissection, dissociation, FACS sorting of single tdTomato^+^ spiral ganglion neurons for a bulk of 100 cells RNAseq analysis. **(b)** The volcano plot shows the change in protein-coding gene expression levels in the *Isl1CKO* compared to control spiral ganglion neurons (adjusted p-value < 0.05, and fold change > 2 cutoff values). The complete list of identified 332 down- and 318 up-differentially expressed genes is in Supplementary Data 1. **(c)** Enrichment map of down- and upregulated gene ontology (GO) sets visualized by the network. Each node represents a GO term; edges depict shared genes between nodes. Node size represents a number of genes of the mouse genome per the GO term, and node fill color represents a GO term significance. Each GO set cluster was assigned with representative keywords; a list of GO sets is available in Supplementary Data 2).

In contrast, upregulated genes particularly enriched in *Isl1CKO* neurons were associated with “regulation of synapse organization, synapse structure or activity” and “regulation of synapse assembly”, indicating compensatory changes relating to the ability of neurons to form neuronal projections towards their targets. Upregulated genes encoding molecules critical for synapse formation and adhesion included several members of the cadherin superfamily (*Cdh7, Cdh10, Pcdh8, Pcdh9, Pcdh10, Pcdh18*, and *Pcdh19*), adhesion-related genes (*Ptprd, Ptprs, Cntnap2, Itga4, Itga5*), and the ephrin ligands (*Efna3, Efna5*). Interestingly, “synaptic signaling” was among the GO terms with significant representation in both up- and down-regulated genes, suggesting changes in synaptic circuits and neuronal activity. We also found many genes encoding transcription factors and signaling molecules, indicating that ISL1 may act through transcription networks instead of defined target genes. The expression of neural-specific bHLH factors important for differentiation and maturation of neurons was increased in *Isl1CKO*, including members of NeuroD (*Neurod6, Neurod2*) Nscl (*Nhlh2*), and Olig (*Bhlhe23*) families ^30^. Genes encoding LIM domain proteins, the transcription factors *Lhx1, Lhx2, Lmo2, Lmo3*, a core member of the PCP signaling paradigm, *Prickle1*, and neuronal homeobox genes, *Emx2, Uncx*, and *Pknox2* were upregulated in *Isl1CKO* neurons. *Prickle1* is an essential regulator of neuron migration and neurons’ distal and central projections in the cochlea ^31^. Some of the identified genes encoding regulatory molecules shown to be essential for neuronal development, the formation of SGNs, and their projections were downregulated in *ISL1CKO* neurons, such as the signaling molecules *Shh* ^32^, *Wnt3* ^3^, and *Fgfs* (*Fgf10, Fgf11*, and *Fgf13*) ^33, 34^, and the transcription factors *Gata3* ^12, 13^, *Irx1, Irx2, Pou3f2, Pou4f2* ^22^, and *Runx1* ^3, 4^.

Consistent with the idea that ISL1 may control a gene expression program mediating different aspects of neuronal development in the cochlea, several identified genes are known to affect the formation of SGNs and their innervation patterns. For instance, a mouse model with EphA4 deficient signaling in SGNs results in aberrant mistargeting of type I projections to OHCs ^35^ and SGN fasciculation defects ^36^. Targeted disruptions of the neurotrophin receptors, *Ntrk2* and *Ntrk3*, are associated with misplaced SGNs in the modiolus, disorganized projections, and reduced neuronal survival ^37^. The neurotrophin receptor p75NTR (*Ngfr*)-deficient SGNs demonstrate enhanced neurite growth behavior ^38^. Mutation of *Prickle1* results in disorganized apical afferents ^31^. Reduced and aberrant cochlear innervation was shown in mice with *Gata3*-specific SGN deletion ^13^. *Nrp2* expressed in SGNs is required for axon path finding and a normal pattern of cochlear innervation ^39^. The axon guidance molecule SLIT2 and its receptor ROBO1/2 control precise innervation patterns, and their deletions induce misplacement of SGNs ^40^. Mice lacking *Dcc* have the more severe phenotype of mislocated SGNs and disorganized central projections, although most SGNs of *Dcc^-/-^* remained within Rosenthal’s canal ^41^.

Other significantly deregulated genes mediate several aspects of neural-circuit formation in the brain, including neurite elongation, axon guidance, and interactions between neurons and microglia. For instance, deletion of *Epha5* alters retinal axon guidance within the midbrain ^42^; inhibition of *Ntng1* reduces microglial accumulation and loss of cortical neurons ^43^; the absence of either *Bhlhb22* or *Prdm8* alters neural circuit assembly ^*44*^; LHX1 controls neuronal migration and axonal projections of motor neurons ^*45*^; and LHX2 influences axon guidance and the topographical sorting of axons by regulating ROBO1/2 ^46^. These molecular differences dovetail well with abnormalities in the innervation pattern and SGN migration defects *Isl1CKO.* Selected differentially expressed genes from the RNAseq analysis, namely *Dcc, Epha5, Ntng1, Prdm8, Robo2, Slit2, Tbx3, Uncx, Gata3, Lhx1, Lhx2, Cdh7, Nhlh2, Ntrk2*, and *Ntrk3*, were further validated by RT-qPCR of RNA isolated from whole inner ears of E14.5 embryos (Supplementary Fig. 4). These changes indicate that ISL1 regulates transcriptional networks that underlie neuronal identity and function during differentiation of SGNs and that *Isl1* elimination results in a major impairment in the development, axonogenesis, migration, and molecular characteristics of these neurons.

### *Isl1CKO* mice have altered hearing function

Considering the substantial abnormalities in the formation of the spiral ganglion, innervation, and molecular neuron features, we assessed the hearing function of *Isl1CKO* mice. We evaluated distortion product otoacoustic emissions (DPOAE) to determine the robustness of OHC function and cochlear amplification. Otoacoustic emissions are a physiological byproduct of an active amplification mechanism when sound-induced vibrations are amplified in a frequency-specific manner by the OHCs of the organ of Corti ^47, 48^. Compared to controls, the DPOAE responses of *Isl1CKO* were significantly reduced in the frequency range between 4 and 24 kHz (Fig. 4a). DPOAE amplitudes at frequencies of 26 kHz and higher were comparable between control and *Isl1CKO* mice. Based on the physiological place-frequency map in the normal mouse cochlea ^49^, frequencies above 26 kHz are located at the basal half of the cochlea. Thus, decreased DPOAE responses may be attributed to the more profound morphological abnormalities in the apex, disorganized innervation, and disorganized rows of OHCs in the apical end.

**Figure 4.**
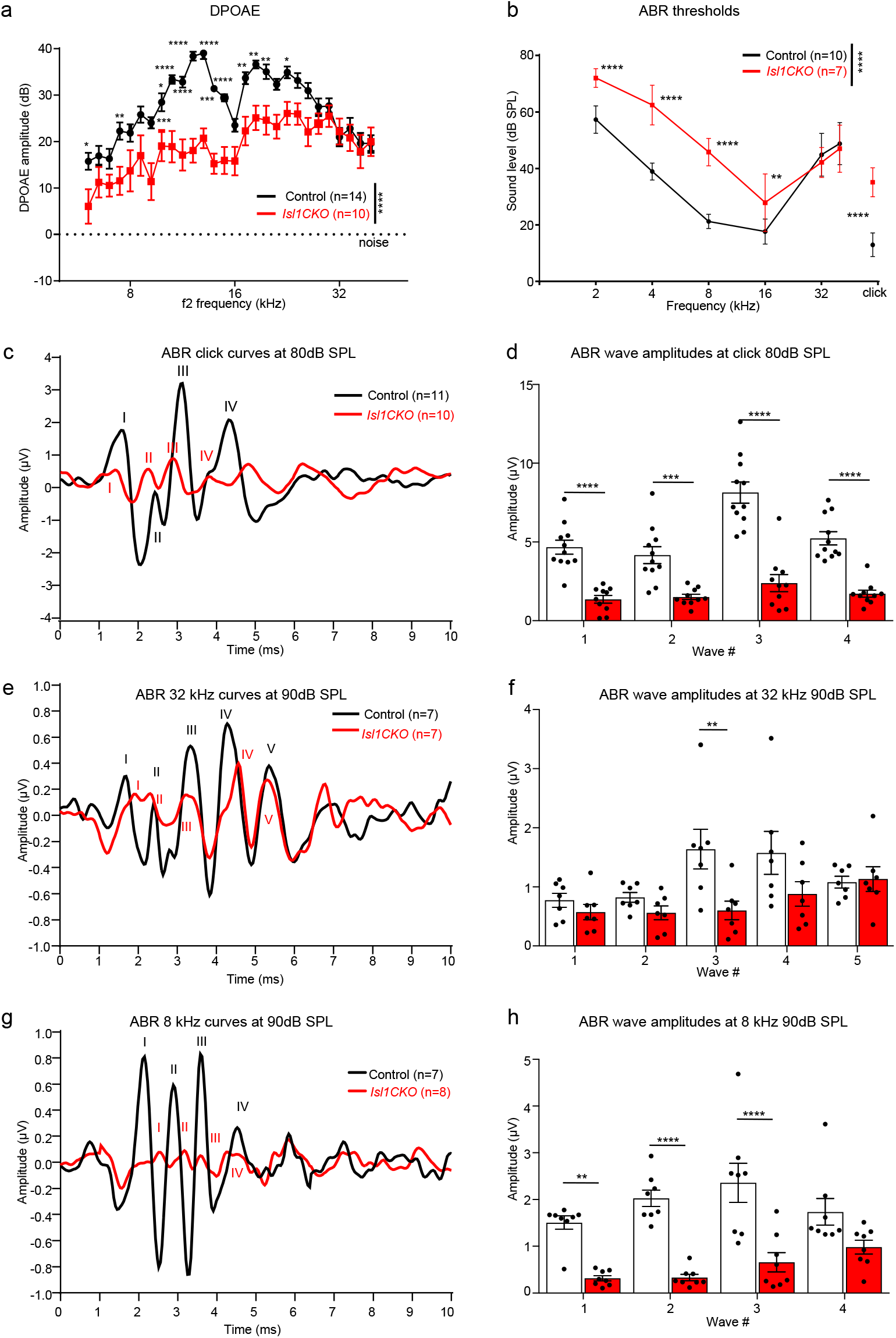
Hearing impairment is detected in *Isl1CKO* mice. (**a**) Distortion product otoacoustic emissions (DPOAEs) show significantly reduced levels in the low and middle-frequency range (4-24 kHz). Data are expressed as mean ± SEM, two-way ANOVA with Bonferroni post-hoc test, *P < 0.05, **P < 0.01, ***P < 0.001 ****P < 0.0001. (**b**) The average auditory brainstem response (ABR) thresholds of *Isl1CKO* and control mice are analyzed by click-evoked ABR. Data are expressed as mean ± SD, two-way ANOVA with Bonferroni post-hoc test, **P < 0.01, ****P < 0.0001. **(c)** Averaged ABR response curves evoked by an 80 dB SPL click; (**e**) by a 90 dB SPL pure tone of 32 kHz frequency; and (**g**) by a 90 dB SPL pure tone of 8 kHz frequency are presented. (**d, f, h**) Averaged individual ABR wave amplitudes are shown for the corresponding peaks. Data are expressed as mean ± SEM, two-way ANOVA with Bonferroni post-hoc test, **P < 0.01 ***P < 0.001, ****P < 0.0001.

We evaluated auditory brainstem responses (ABRs), which measure electrical activity associated with the propagation of acoustic information through auditory nerve fibers. Measurements of ABR thresholds showed that all *Isl1CKO* animals displayed elevated thresholds indicative of hearing loss compared to age-matched control animals, except at frequencies 32 and 40 kHz, which were comparable to ABR thresholds in control mice (Fig. 4b). Using click-evoked ABR, we evaluated waveform characteristics associated with the propagation of acoustic information through the auditory nerve to higher auditory centers (Fig. 4c). Wave I reflects the synchronous firing of the auditory nerve. In contrast, waves II–V are attributed to the electrical activity of downstream circuits in the CN, superior olivary complex, lateral lemniscus, and inferior colliculus ^50^. Waves IV and V are products of the sound-evoked neuronal activity in the lateral lemniscus and inferior colliculus ^51, 52, 53, 54, 55^. The waves IV and V are often paired and superimposed in commonly used non-transgenic mouse strains ^52, 55^. The ABR traces in all our panels were averaged traces from all measured animals. A part of the animals in both experimental groups had a hardly distinguishable wave V in response to the click and 8 kHz sound stimulation; therefore, to prevent bias, we did not include wave V. The amplitudes of ABR waves I-IV were significantly reduced in *Isl1CKO* (Fig. 4d). Since the ABR threshold for 32 kHz and above were comparable between age-matched controls and *Isl1CKO* mutants (Fig. 4b), we used the pure-tone stimuli of 32 kHz to evaluate ABR responses (Fig. 4e). A significant difference amongst the genotypes was found only for amplitude reduction of wave III (Fig. 4f), thus indicating preserved synchronized activities of peripheral and brainstem auditory processing. Although waves I and II amplitude for both mutant and control mice were similar, there were apparent differences in the ABR waveform morphology. The latency of ABR wave I was delayed, the relative interwave latency between peaks I and II was shortened, and the trough between wave I and II diminished, resulting in a fusion of both peaks in *Isl1CKO*. A delay of the leading peak of ABR wave I recovered towards ABR wave III. The wave I and II changes reflect abnormalities in the summated response from SGNs, auditory nerve fibers, and most likely the CN.

Additionally, we used a pure-tone stimulus of 8 kHz to evaluate ABR responses (Fig. 4g), as ABR thresholds for 8 kHz were significantly reduced for *Isl1CKO* mice. In contrast to ABR amplitudes at 32 kHz stimuli, ABR amplitudes for 8 kHz pure-tone stimuli were significantly reduced for I-III waves (Fig. 4h). The results indicate abnormalities in the cochlear auditory neurons and propagation of acoustic information through auditory nerve fibers to higher auditory centers. We observed a marked drop in the wave I growth function. Still, by comparing wave I and IV peaks, an increased central gain was noted in *Isl1CKO* (Supplementary Fig. 5), indicating compensatory plasticity at higher auditory circuits for cochlear damage with diminished afferent input ^56^.

### *Isl1CKO* mice display abnormalities in the ascending auditory pathways

Having recognized uneven hearing function loss, as measured by ABRs, we further wanted to investigate the distribution of SGNs in the *Isl1CKO* cochlea and the morphology of the components of the central auditory pathway. We generated 3D visualization of the cochlea using light-sheet fluorescent microscopy (Fig. 5a-i, Supplementary Videos S1-S4). In the first cochlear preparation, neurons were visualized by *tdTomato* reporter expression and hair cells in the organ of Corti were labeled using anti-Myo7a. In the second preparation, tdTomato^+^ neurons were co-labeled with anti-NeuN, a nuclear marker of differentiated neurons. Compared to the recognizable coil of cochlear neurons of the spiral ganglion in the controls (Fig. 5a-c, Supplementary Videos S1, S2), SGNs were aberrantly distributed in the *Isl1CKO* cochlea (Fig. 5d-i, Supplementary Videos S3, S4). Many *Isl1CKO* neurons were in the center of the cochlea, the modiolus, that normally contains only central processes of SGNs, as shown in the control cochlea (Fig. 5b, c). Some *Isl1CKO* neurons were found in their proper position in the Rosenthal’s canal, forming a dense strip of neurons, suggesting a partially preserved organization of the spiral ganglion (Fig. 5f, i). This segment of the spiral ganglion formed in the mid-base of the *Isl1CKO* cochlea correlates with the estimated characteristic place in the cochlear place-frequency map for the characteristic frequency of 32 kHz ^49^ (Fig. 5j). A partial preservation of the spiral ganglion in the mid-base corresponds to the retained high-frequency hearing function of *Isl1CKO*. Identical to the SGN migration abnormalities recognized at P0, most SGNs were in the modiolus, outside of Rosenthal’s canal, in *Isl1CKO* adult mice (Supplementary Fig. 6). Apart from the changes in the cochlear ganglion, the vestibular ganglion was fused and enlarged in *Isl1CKO* compared to distinct superior and inferior vestibular ganglia in controls (Fig. 5k, l). The spatial segregation between the vestibular and spiral ganglion was diminished in *Isl1CKO*, with some areas fused, and thus, forming an aberrant “spiro-vestibular” ganglion.

**Figure 5.**
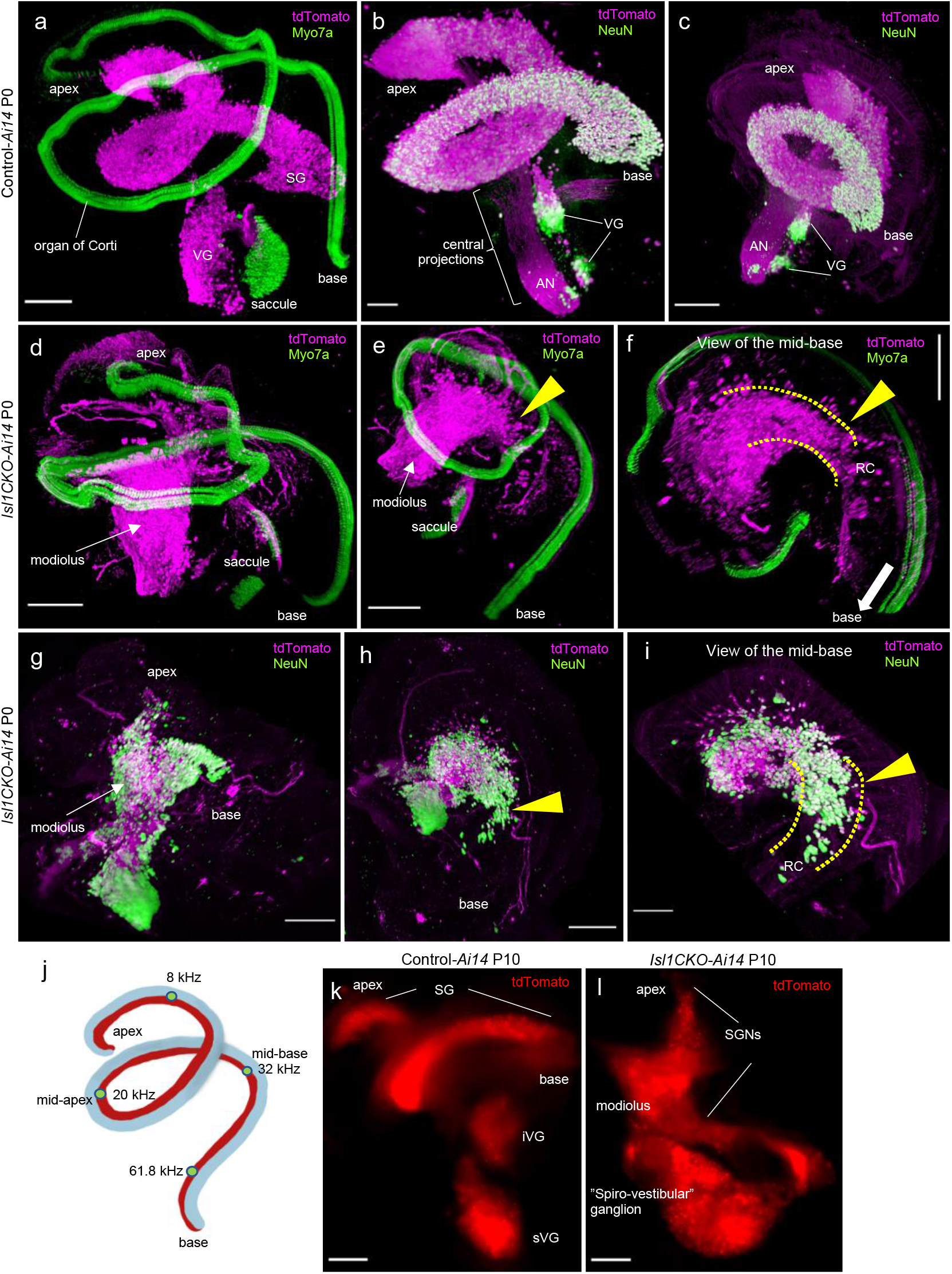
Neurons in the *Isl1CKO* inner ear are abnormally distributed. **(a-i)** Microdissected cochleae of reporter control-*Ai14* and *Isl1CKO-Ai14* mice were cleared (CUBIC protocol), immunolabeled, imaged, and reconstructed in 3D using light-sheet fluorescence microscopy (see Supplementary Videos S1-S4). (**a**) In the control cochlea, tdTomato^+^ SGNs form a coil of the spiral ganglion (SG) in the Rosenthal’s canal and anti-Myo7a labeled hair cells show the spiral shape of the organ of Corti. Parts of the vestibular ganglion (VG) with tdTomato^+^ neurons and the saccule with Myo7a^+^ hair cells are shown. (**b, c**) Anti-NeuN (a nuclear marker of differentiated neurons) co-labeled tdTomato^+^ neurons form SG and VG in the control cochlea; the modiolus contains the central processes, forming the auditory nerve (AN). (**d-f**) In *Isl1CKO-Ai14*, tdTomato^+^ neurons are mainly located in the conical central part of the cochlea, the modiolus, in contrast to the spiral of the organ of Corti with Myo7a^+^ hair cells. Some *Isl1CKO* neurons form a segment of the spiral ganglion in the Rosenthal’s canal (RC) in the mid-base region (arrowhead), shown in detail (**f**), the RC area is highlighted by dotted lines. (**g-h**) Abnormal and uneven distribution of cochlear tdTomato^+^ neurons is shown by anti-NeuN co-labeling. Many neurons are misplaced in the modiolus, but some neurons are in their close-normal position in RC (arrowhead). **(j)** Schematic drawing of the mouse cochlea shows the cochlear place-frequency map for four cochlear regions termed the apex (corresponding to the best encoding frequencies ~8 kHz at the distance from the basal end of 82 %), mid-apex (~19 kHz at 49% length), mid-base (~32 kHz at 32% length), and basal turn (~62 kHz at 9% length) ^49^. **(k)** Whole mount images of the P10 inner ear show distribution of tdTomato labeled neurons in superior (sVG) and inferior vestibular ganglia (iVG), and SG in the control-*Ai14*. **(l)** In the *Isl1CKO-Ai14* mutant, the VG is fused and enlarged, spatial segregation between VG and SG is diminished, forming an aberrant “spiro-vestibular” ganglion, and SG has lost its spiral shape. SGNs, spiral ganglion neurons. Scale bars: 200 μm, except for 100 μm (b, i).

Using dye tracing, we evaluated the segregation of central axons of the auditory nerve (the cranial nerve VIII; Fig. 6a, schematic view of dye applications). In controls, the central axons labeled by dyes applied into the base and apex are segregated in the auditory nerve and from the vestibular ganglion, labeled by dye injected into the vestibular end-organs (Fig. 6b, Supplementary Fig. 7). In contrast, the central axons from the cochlear base and apex virtually overlapped in the auditory nerve (yellow fibers), and neurons labeled by cochlear dye applications were detected to be intermingled with vestibular neurons in an aberrant “spiro-vestibular” ganglion in *Isl1CKO* (Fig. 6c). Unfortunately, the mixing of spiral and vestibular ganglion neurons in *Isl1CKO* mice precluded a full quantitative assessment. The unusual fibers labeled by the apical dye application were looping around the vestibular ganglion in *Isl1CKO*.

**Figure 6.**
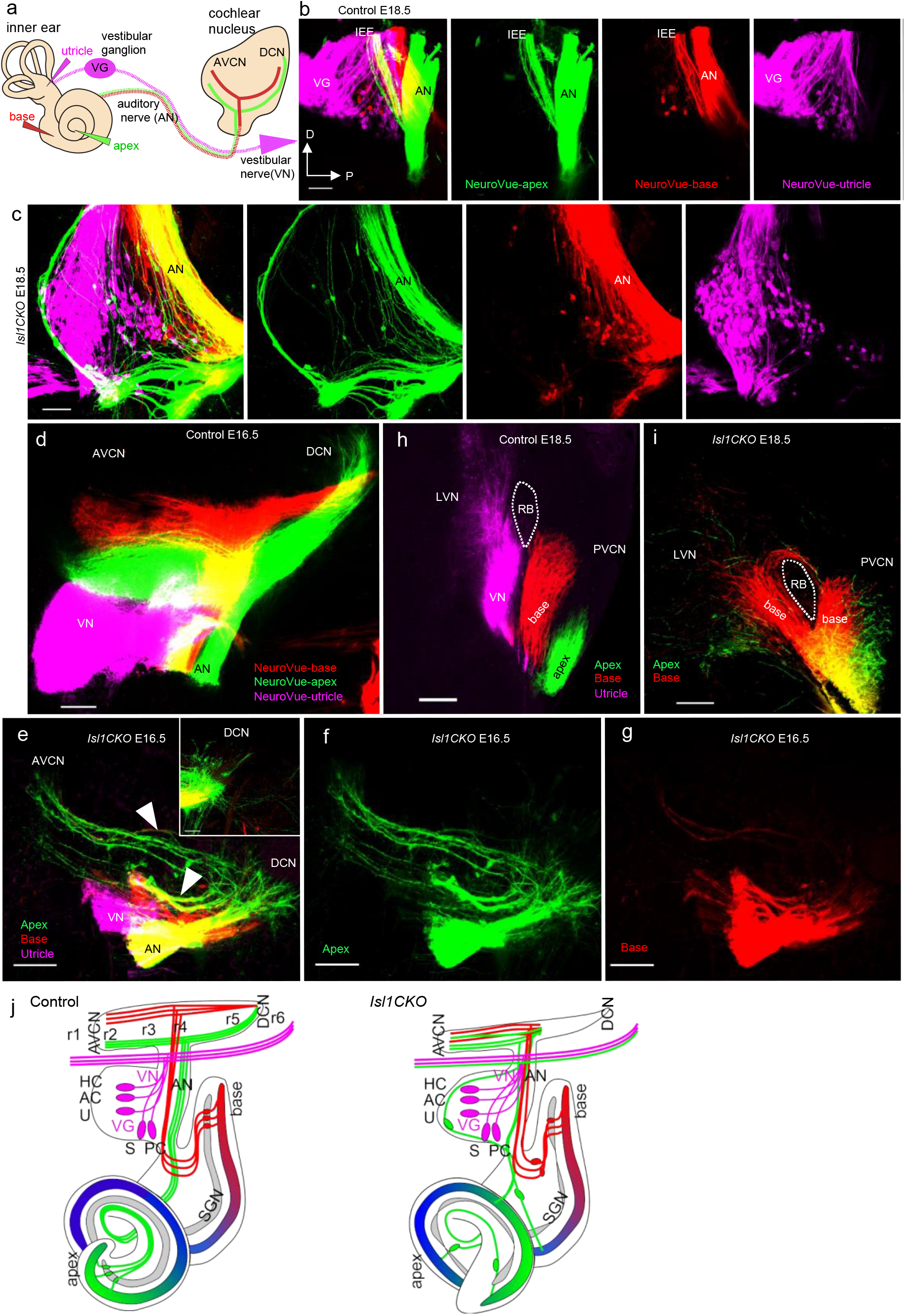
*Isl1CKO* neurons project unsegregated and disorganized central projections to the cochlear nucleus. **(a)** The schematic diagram visualizes dye tracing from the inner ear and its connections to the auditory brainstem, using insertions of differently colored NeuroVue dyes into the vestibular end-organ (utricle, magenta), cochlear base (red), and apex (green). Axonal projections from cochlear neurons to the cochlear nucleus (CN) bifurcate with one branch synapsing in the dorsal (DCN) and the other innervating the anteroventral CN (AVCN). **(b)** Images of individual colors of the separate channels and a merged image show distinct and spatially restricted bundles of neuronal fibers of the auditory nerve (AN) projecting to the CN and the vestibular ganglion (VG) in controls, labeled by dye applications into the apex (green), base (red), and utricle (magenta). The only mixed bundle (yellow) is inner ear efferents (IEE). **(c)** In *Isl1CKO*, the segregation of central axons is lost, as fibers labeled from the apex (green) and base (red) are mainly overlapping in the AN (yellow fibers), and neurons labeled by dyes applied into the cochlear base and apex are mixed with VG neurons to form an aberrant ganglion, the “spiro-vestibular” ganglion. Note that fibers labeled from the apex (green) form an unusual fiber loop around the ganglion, and no IEE are recognizable in *Isl1CKO*. **(d)** The tonotopic organization of the CN subdivisions is shown with low-frequency afferents labeled from the apex (green) and high frequency from the base (red) and organized as parallel fibers in isofrequency bands in controls at E16.5. **(e)** In *Isl1CKO*, cochlear afferents enter AVCN as a single bundle instead of forming separate basal and apical projections. The branch synapsing in the AVCN is reduced and disorganized. Arrowheads indicate overlapping fibers. The DCN branch is represented by just a few fibers in *Isl1CKO* (inset in **e**). **(f, g)** Images of individual colors of the separate channels represent projections labeled from the apex (green) and base (red), shown as a merged image in e. **(h, i)** The confocal section shows segregated cochlear afferents from the base and apex, and vestibular afferents extend toward the control’s lateral vestibular nucleus (LVN). In contrast, cochlear afferents of *Isl1CKO* overlap (yellow fibers), and some basal afferents extend toward the CN, and some project toward the LVN and loop back to the CN. (**j**) Graphic summary of aberrant location and projections of auditory neurons in *Isl1CKO*. In the control, the auditory system is tonotopically organized, including neurons of the spiral ganglion, the auditory nerve, and the CN subdivisions. In *Isl1CKO*, excluding some portion of neurons in the base, neurons are located outside of Rosenthal’s canal. Some neurons projecting to the cochlea are mixed with vestibular neurons. The central axons from the apex and base are not segregated in the AN and AVCN. Projections entering the DCN are diminished in *Isl1CKO*. AC, anterior crista; D, dorsal; HC, horizontal crista; P, posterior axes; PVCN, posteroventral CN; PC, posterior crista; RB, restiform body; r1-r6, rhombomere 1 - 6; U, utricle; S, saccule; VN, vestibular nerve. Scale bars: 100 μm.

The CN is the first structure of the ascending auditory pathways, where the auditory nerve fibers project. The auditory nerve bifurcated with one branch, synapsing in the posteroventral (PVCN) and dorsal (DCN) CN and the other innervating the anteroventral CN (AVCN; Fig. 6a). Dye tracing showed segregated projections of apical and basal cochlear afferents forming parallel isofrequency bands in controls at E16.5 (Fig. 6d). In contrast, comparable injections in *Isl1CKO* showed that axonal projections to the CN were reduced, restricted, disorganized, and lacked clear apex- and base-projection segregations, as many fibers overlapped (Fig. 6e-g). Only a few fibers can occasionally be seen expanding to the DCN in *Isl1CKO* at E16.5 (inset in Fig. 6e). Aberrant central projections were further demonstrated by cochlear neurons projecting to the lateral vestibular nucleus (LVN) in *Isl1CKO* (Fig. 6i). No such projections were found in control mice with segregated vestibular projections to the vestibular nucleus, and basal and apical afferents reaching the CN (Fig. 6h). Collectively, these data show disorganized central projections and a loss of tonotopic organization of both the auditory nerve and the CN in *Isl1CKO*, as the apical and basal projections are not completely segregated (see graphic summary in Fig. 6j).

Since the size and number of neurons in the CN depend on input from the auditory nerve during a critical development period up to P9 ^57^, we also analyzed the volume of the CN of *Isl1CKO*. The volume of the CN of adult mutants was reduced by approximately 50% compared to controls at postnatal day P35 (Fig. 7a, b, g). As *Isl1* is not expressed in the CN ^19^, the reduced size is likely a secondary effect associated with reduced afferent input consistent with the previously reported impact of neonatal cochlear ablation ^2^. The spherical and globular bushy cells are principal cells that receive large auditory nerve endings, called “endbulbs of Held” and “modified endbulbs”, specialized for precise temporal firing ^1, 58^. Using anti-parvalbumin to label spherical bushy cells and anti-VGlut1 to label auditory-nerve endbulbs of Held ^59, 60^, we demonstrated that auditory afferents of *Isl1CKO* and controls formed comparable clusters of boutons that wrap the somas of their targets (Fig. 7c-f). Although the CN of *Isl1CKO* was smaller, the density of bushy cells (number of parvalbumin^+^ neurons per μm^2^) was similar in controls and *Isl1CKO* (Fig. 7h).

**Figure 7.**
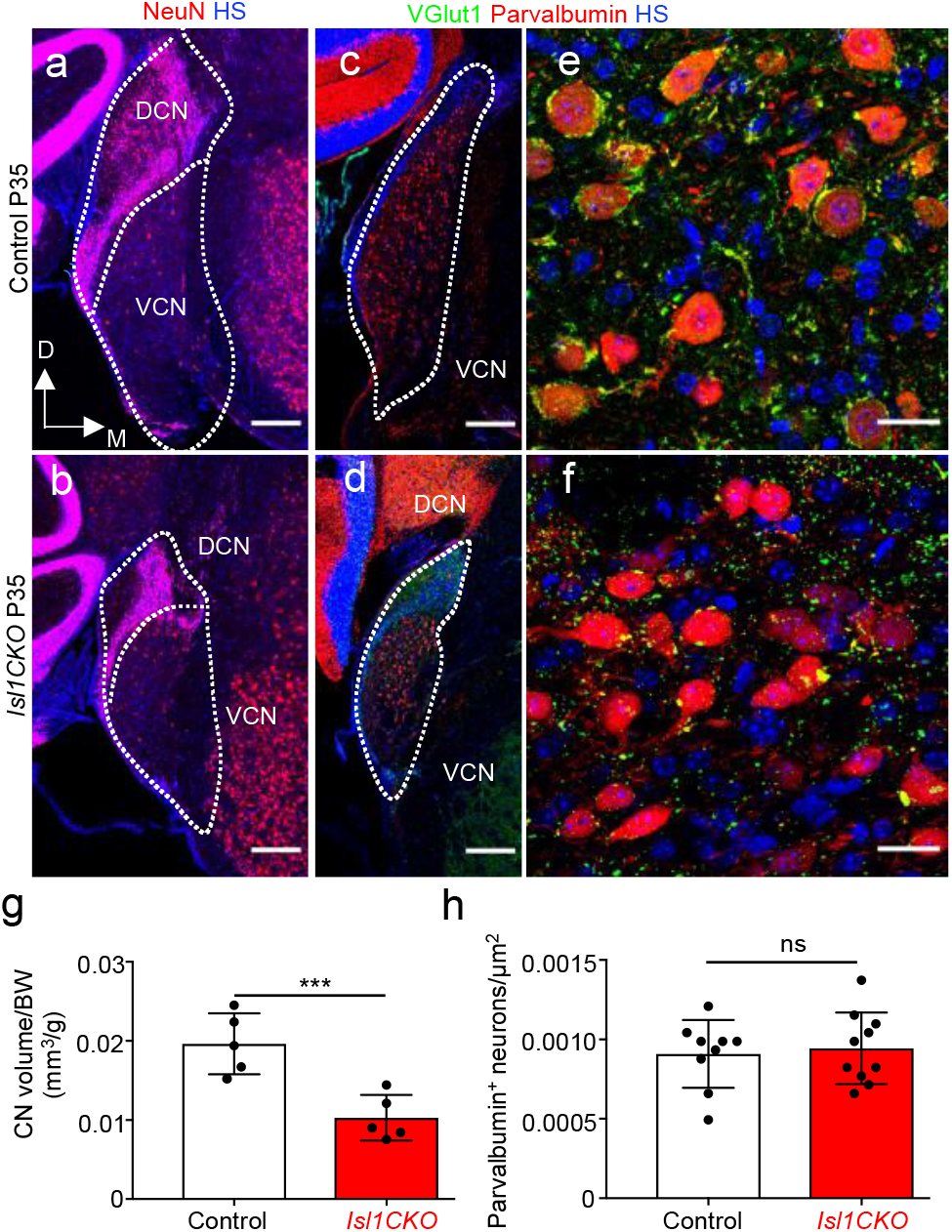
The cochlear nucleus of *Isl1CKO* is reduced. **(a, b)** Coronal sections of a brain (immunostained with anti-NeuN) of adult controls and *Isl1CKO*, show the DCN and VCN; the dotted line indicates the boundaries of the CN. **(c-f)** Representative images of immunolabeled brain sections with anti-parvalbumin to label the sphe3ri6c0al bushy cell soma and anti-VGlut1 to label auditory-nerve endbulbs of Held around the bushy cells. Higher-magnification images show the distribution of parvalbumin^+^ neurons and the presence of VGlut1^+^ auditory nerve synaptic terminals. (**g**) Quantification of the adult CN volume, adjusted to body weight (n = 5 mice per genotype) and **(h)** parvalbumin^+^ neurons per μm^2^ of the AVCN (n = 3 mice per genotype, 2-4 sections per mouse). Data are expressed as mean ±SD, unpaired *t-test* (***P < 0.001; ns, not significant). D, dorsal; M, medial axis; HS, Hoechst nuclear staining; DCN, dorsal cochlear nucleus; VCN, ventral coch3l7ea1r nucleus. Scale bars: 200 μm (a-d); 20 μm (e,f).

### Characteristics of inferior colliculus neurons are distorted in *Isl1CKO*

We evaluated inferior colliculus (IC) properties, which is the principal auditory structure of the midbrain for the ascending auditory pathways and descending inputs from the auditory cortex ^61^. The IC allows for sound localization, integrates multisensory and non-auditory contributions to hearing, and plays an essential role in generating the startle response. We demonstrated no significant IC size reduction in *Isl1CKO* compared to controls (Fig. 8a, b). We compared neuronal characteristics in the central nucleus of the IC of *Isl1CKO* and control animals using multichannel electrodes. Extracellular electrophysiological recordings of neuronal activity in controls showed a well-defined narrow single-peaked profile of the excitatory receptive fields (Fig. 8c). Compared to simple narrow V-shape receptive fields in controls, we recorded the multipeaked broad tuning curves in the IC of *Isl1CKO* mice, suggesting multiple inputs from the lower levels of the auditory system and overall, the worsened tunning characteristics of IC neurons.

**Figure 8.**
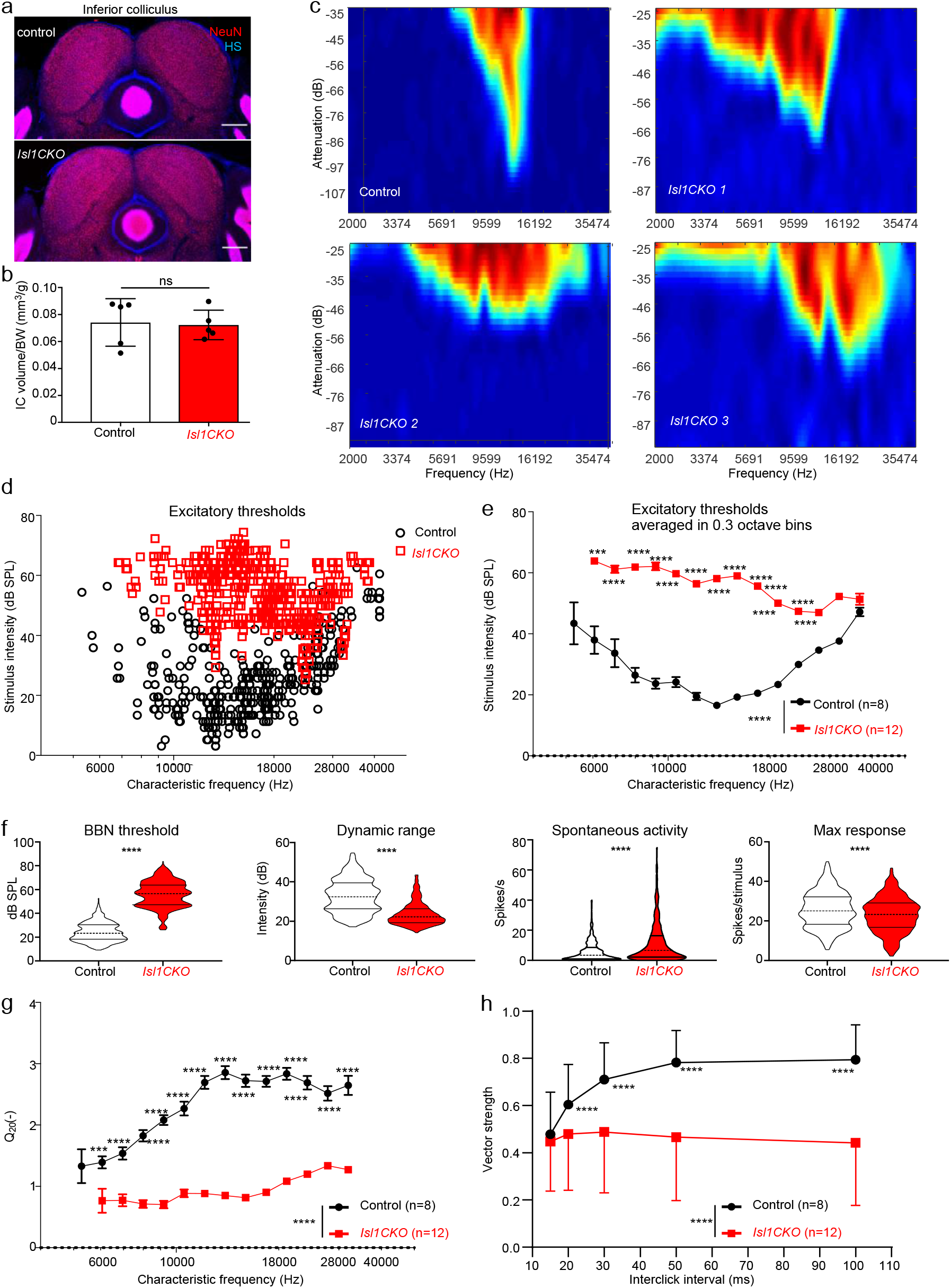
Characteristics of IC neurons (units) are affected in *Isl1CKO* mice. **(a)** Immunostaining of coronal brain sections for NeuN and nuclear staining Hoechst (HS). Scale bars, 200 μm. **(b)** Quantification of the adult control volume and *Isl1CKO* inferior colliculus (IC) adjusted to body weight. Data are expressed as mean ± SD (n = 5 mice per genotype), unpaired *t-test* (ns, not significant). **(c)** Representative examples of tuning curves recorded in the IC display impairments in tuning properties with broad and irregular receptive fields in *Isl1CKO* compared to control mice. **(d)** Scatter diagram shows the distribution of the excitatory thresholds of IC neurons in dependency on the characteristic frequency (CF) recorded in control and *Isl1CKO* mice. Note, no IC neurons are recorded below 6 kHz in *Isl1CKO* mutants. **(e)** Excitatory thresholds of the IC neurons at different CFs are shown as averages in 0.3-octave bins in control and *Isl1CKO* mice. Data are expressed as mean ± SEM. Two-way ANOVA with Bonferroni post hoc test. **P < 0.01, ***P < 0.001, ****P < 0.0001. **(f)** Comparison of the rate intensity function parameters between control (n = 8) and *Isl1CKO* mice (n = 12): broadband noise (BBN) threshold, dynamic range, spontaneous activity, and maximum response magnitude. Violin plots indicate median (middle line), 25th, and 75th percentile (dotted lines), unpaired *t*-test, ****P < 0.0001. **(g)** The sharpness of the neuronal tuning expressed by quality factor Q_20_ (the ratio between the CF and bandwidth at 20 dB above the minimum threshold) averaged in 0.3-octave bins is decreased in *Isl1CKO*. **(h)** Synchronization of units with click trains. Vector strength computed for different inter-click intervals. Data are expressed as mean ± SD, two-way ANOVA with Bonferroni post-hoc test, ****P < 0.0001.

The investigation of the responsiveness of IC neurons (IC units) to different sound frequencies showed higher excitatory thresholds in *Isl1CKO* compared to control animals in measured frequencies between 6 to 28 kHz (Fig. 8d, e). However, responsiveness of IC units to recorded frequencies above 28 kHz was comparable between control and mutant mice. These comparable thresholds for high frequencies between control and *Isl1CKO* mice are consistent with the ABR measurements. At the low frequency range, below 6 kHz, we did not record any IC neurons in *Isl1CKO*. Compared to the controls, the IC neuronal responses in *Isl1CKO* had a higher broadband noise (BBN) threshold, a narrower dynamic range, higher spontaneous activity, and a lower maximum response magnitude (Fig. 8f). The results suggest a functional reduction in sensitivity to sound, audibility, intensity discrimination, and increased excitability of IC neurons in *Isl1CKO*. A commonly used metric unit of auditory tuning is the “quality factor”, or Q, defined as the characteristic frequency (CF) divided by the bandwidth, measured at 20 dB above the minimum threshold (Q_20_). Results revealed a significantly lower quality factor in the mutant mice (Fig. 8g), showing substantially worsened frequency selectivity.

To evaluate the precise temporal representation of sound into the central auditory system, we performed an acoustic stimulation of the IC units with trains of five clicks with different inter-click intervals from 100 ms up to 15 ms. In control mice, increasing time interval between clicks led to a better synchronization of neuronal responses with the individual clicks in the train, implying precision and reliability of temporal sound discrimination ability (Fig. 8h). In contrast, the precise temporal decoding in *Isl1CKO* was disrupted, as the synchronization of neuronal responses was significantly lower for the whole range of inter-click intervals. In the case of the *Isl1CKO*, the synchronization level of neuronal responses remains almost constant, suggesting a lack of precise temporal sound processing.

### The auditory behavior of *Isl1CKO* mice is altered

Next, we evaluated the behavioral responses of *Isl1CKO* mice to sound stimuli. Since the vestibular dysfunctions might influence auditory behavioral responses, we first assessed the vestibular and motor function of *Isl1CKO*. During open-field observations, *Isl1CKO* mice did not display any signs of abnormal locomotor behavior, such as ataxia, difficulty maintaining balance, wagging or vertical bobbing movements of the head, or circling movements. The air-righting test, a basic vestibular function test, and motor function tests on the rotarod demonstrated comparable motor coordination between control and *Isl1CKO* mice (Supplementary Fig. 8). Concurrently, the size of the sensory epithelia of the vestibular organs and the size of the dorsal root ganglia were unaffected in *Isl1CKO* (Supplementary Fig. 8). Thus, locomotor activities of *Isl1CKO* were comparable to control mice. The acoustic startle response (ASR) is usually used as a behavioral readout of hearing status mediated by a brainstem circuit linking cochlear root neurons to spinal motoneurons. The structural basis of the ASR includes cochlear root neurons, CN neurons, the nucleus of the lateral lemniscus, the caudal pontine reticular nucleus, spinal interneurons, and spinal motor neurons ^62, 63, 64^. Similar to the ABR thresholds, the ASR thresholds of *Isl1CKO* significantly increased for startle tone stimuli of 8 kHz and BBN, but not for the high-frequency startle tones (Fig. 9a). The peak latency of the ASR to the BBN stimulation at the 110 dB SPL intensity was prolonged in *Isl1CKO* (Fig. 9b), indicating a slower reaction to the acoustic stimuli. We found significantly reduced ASR amplitudes for all tested sound stimuli at higher intensities, showing deteriorated acoustic startle reactivity in *Isl1CKO* mice (Fig. 9c-e).

**Figure 9.**
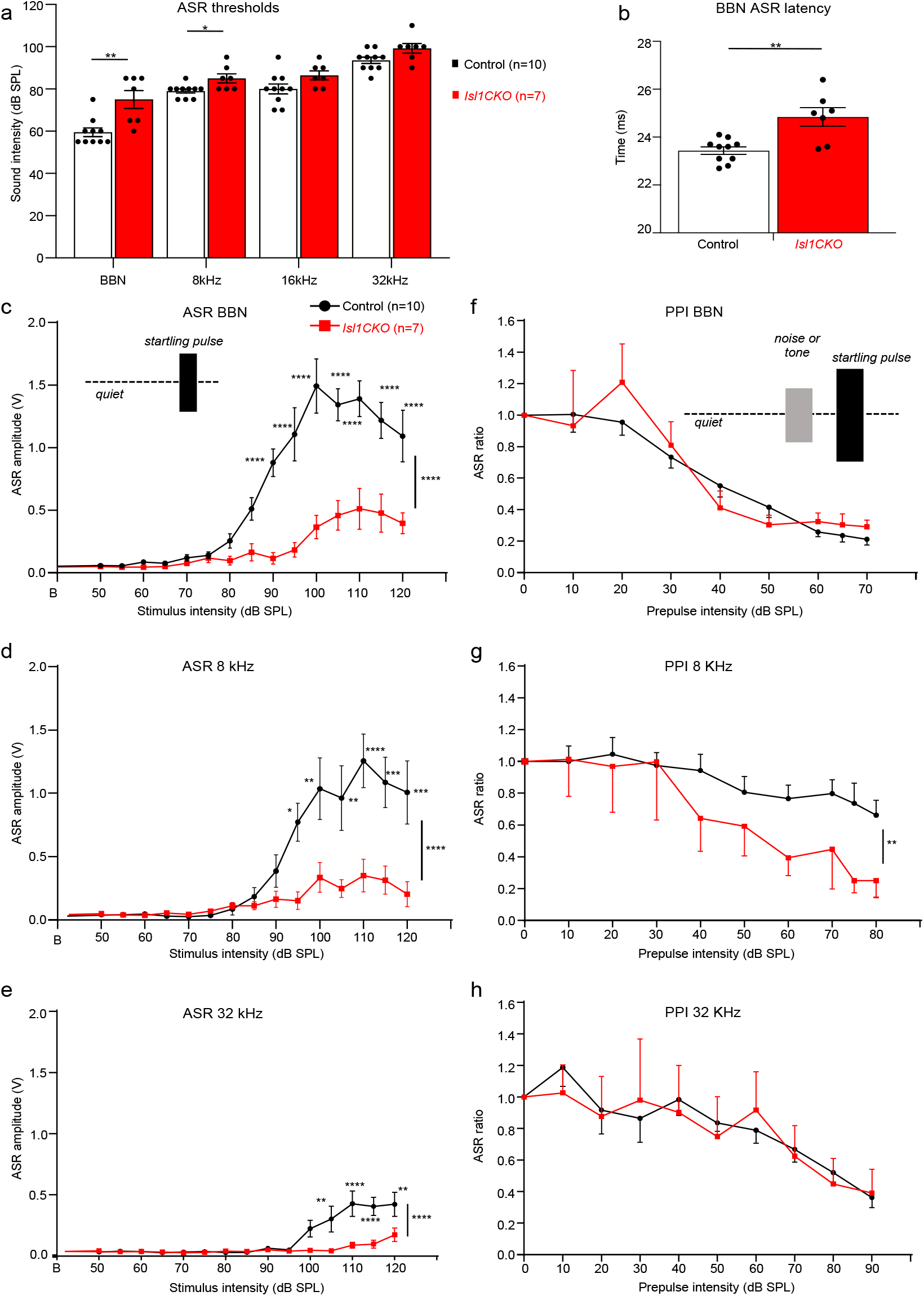
The acoustic startle reflex (ASR) and prepulse inhibition (PPI) responses are altered in the *Isl1CKO* mutant. **(a)** ASR thresholds for broadband noise (BBN) bursts and tone pips at 8, 16, and 32 kHz in control and *Isl1CKO* mice are shown. Holm-Sidak method multiple comparison *t*-test. *P < 0.05, **P < 0.01. **(b)** Significantly increased ASR latency to BBN is found in *Isl1CKO* compared to control mice; unpaired *t*-test, **P < 0.01. **(c)** Amplitude-intensity ASR functions for BBN stimulation and **(d)** for tone pips of 8 kHz and **(e)** 32 kHz at different dB SPL intensities in control and *Isl1CKO* mice are shown. **(f)** Efficacy of the BBN, (**g**) 8 kHz, and (**h**) 32 kHz tone prepulse intensity on the relative ASR amplitudes are displayed; ASR ratio = 1 corresponds to the ASR amplitude without a prepulse (uninhibited ASR). Two-way ANOVA with Bonferroni post hoc tests. *P < 0.05, **P < 0.01, ***P < 0.001, ****P < 0.0001. All data are expressed as mean ± SEM.

To further assess complex auditory discrimination behavior, we exposed control and *Isl1CKO* mice to a prepulse inhibition (PPI) paradigm, i.e., the inhibition of the ASR induced by presenting an acoustic stimulus shortly preceding the presentation of an acoustic stimulus, the startling sound. The circuit mediating a prepulse on the startle reflex involves central structures of the auditory pathway, including the IC and the auditory cortex ^65, 66^. We used either BBN or pure tone pips of 8 and 32 kHz at increasing intensities as a non-startling acoustic stimulus (prepulse) that preceded the startle stimulus in a quiet background. The PPI with the prepulse of 32 kHz, the well-preserved audible frequency in *Isl1CKO*, was comparable between control and mutant mice (Fig. 9h). Interestingly, the prepulse of 8 kHz resulted in a more prominent inhibition of the startle response in *Isl1CKO* than in controls (Fig. 9g), despite the significant hearing deficiency at 8 kHz, as shown by ABR evaluations (Fig. 4). This indicates that the 8-kHz prepulse response was enhanced in *Isl1CKO*, suggesting compensatory neural hyperactivity of the central auditory system ^56, 67^. Thus, the ASR and PPI of startle analyses indicate abnormalities of the acoustic behavior of *Isl1CKO* mutants.

## Discussion

Our study shows for the first time that LIM homeodomain transcription factor ISL1 regulates neuronal development in the cochlea. Using RNA profiling, morphological, and physiological analyses, we provide evidence that ISL1 coordinates genetic networks affecting the molecular characteristics of SGNs, their pathfinding abilities, and auditory information processing. The elimination of *Isl1* in neurons during inner ear development results in a migration defect of SGNs, disorganized innervation in the cochlea, unsegregated and reduced central axons, and reduced size of the CN. This neuronal phenotype of *Isl1CKO* was accompanied by hearing impairment, abnormalities in sound processing in the IC, and aberrant auditory behavior.

ISL1 is critical for the development of multiple tissues, neuronal and non-neuronal cells ^68, 69, 70, 71, 72^. Different aspects of neuronal development depend on ISL1, including specification of motoneurons ^68^, sensory neurons ^25, 70^, axonal growth ^73^, and axonal pathfinding ^74^. During inner ear development, ISL1 is expressed in both neuronal and sensory precursors ^18, 20, 21, 22, 75^. Transgenic modulations of *Isl1* indicate important roles of ISL1 in the maintenance and function of neurons and hair cells and as a possible contributing factor in neurodegeneration ^23, 76, 77, 78^. These studies suggest that ISL1 plays a role in developing neurons and sensory cells, but no direct evaluation of ISL1 function has been performed. To circumvent the pleiotropic effects of *Isl1* in embryonic development, in this study, we used *Neurod1^Cre^* to delete *Isl1* specifically in the inner ear neurons.

We established that ISL1 is necessary for neuronal differentiation programs in the cochlea and the functional properties of the auditory system. Our RNA profiling of SGNs demonstrated transcriptome changes induced by a loss of *Isl1* affecting the molecular characteristics of neurons and pathfinding abilities, including neurotransmission, the structure of synapses, neuron migration, axonogenesis, and expression of crucial guidance molecules. Consistent with the central role of ISL1 in sensory neuron developmental programs ^25^, regulatory networks of signaling molecules and transcription factors were affected in *Isl1CKO* neurons, such as proneural bHLH factors (members of NeuroD, Olig, and Nscl families), LIM-only (*Lmo2, Lmo3*) and LIM homeodomain transcription factors (*Lhx1, Lhx2, Isl2*), transcription activation complexes for coordination of particular differentiation programs Eyes absent (*Eya4* and *Eya2*) and Sine oculis (*Six2*), POU homeodomain trans-regulatory factors (*Pou3f2* and *Pou4f2*), and FGF signaling molecules (*Fgf10, Fgf11, Fgf13, Fgf14*) and their downstream targets (*Etv1, Etv4, Etv5*). Interestingly, the transcription factor *Gata3* was downregulated, suggesting that ISL1 is upstream of the Gata3 transcriptional network of neuronal differentiation programs ^13^. Thus, ISL1 orchestrates a complex gene regulatory network driving multiple aspects of neuronal differentiation in the cochlea and defining neuronal features.

The most striking morphological features of the neuronal phenotype of *Isl1CKO* are the aberrant migration and pathfinding of SGNs. A similar migration deficit was reported for *ErbB2* null mutants ^79^; however, the interpretation of the findings is convoluted by the compounded direct inner ear effects and the effects associated with neural crest-derived Schwann cells. Additionally, conditional deletion of *Sox10* produced by *Wnt1^Cre^* resulted in abnormal migration of SGNs similar to the *Isl1CKO* phenotype ^80^. Migration defects of SGNs in both *ErbB2* ^79^ and *Sox10* mutants ^80^ were attributed to the complete absence of Schwann cells in the entire inner ear ganglion. However, in our *Isl1CKO* mutant, Sox10^+^ Schwann cells were found in a similar density in the spiral ganglion of both mutant and control mice, thus excluding any direct involvement of glial cells in the migration defects of *Isl1*-deficient neurons. We identified several genes with altered molecular profiles that are associated with neuronal phenotypes of projection and migration abnormalities. For instance, misplaced SGNs in the modiolus are in *Ntrk2* and *Ntrk3* neurotrophin receptor mutants ^37^. Less severe phenotypes of misplaced SGNs have been reported for mice lacking *Dcc* ^41^ or mice with deficient Slit/Robo signaling ^40^. All these reported SGN migration defects are also associated with abnormalities in projections. In line with the *Isl1CKO* phenotype of deficient neuronal migration and projections, *Isl1* elimination in SGNs resulted in major transcriptional changes of genes encoding members of all four classes of guidance molecules and their receptors, growth factors, neurotrophic factor receptors, and cell adhesion molecules (NCAM family, integrins, cadherins and protocadherins) among others that are known to participate in molecular networks coordinating neuronal migratory behaviors, axon guidance, and neural circuit assembly.

More profound disorganization of peripheral processes than defects found in *Isl1CKO* or both *ErbB2* null ^79^ and *Sox10* mutants ^80^ was reported for delayed conditional deletion of *Gata3* in SGNs ^13^. Despite severe disorganization of cochlear wiring of this *Gata3* mutant, central projections maintained their overall tonotopic organization within the auditory nerve and the CN ^12, 13^. In contrast, we provide compelling evidence that elimination of *Isl1* in SGNs affected the pathfinding abilities of neurons in the cochlea to form peripheral processes and establish central projections. Such profound disorganization of peripheral and central projections is known for *Neurod1* mutations ^16, 18, 19^. Deletions of *Neurod1* result in miswired, and diminished SGNs, and loss of tonotopy ^16, 19^. Somewhat similar disorganization of central axons with unsegregated auditory nerve fibers, reduced size of the CN, and missing tonotopic organization of synapsing branches in the CN subdivisions was found in our *Isl1CKO*. Although both *Neurod1CKO* and *Isl1CKO* demonstrated significant hearing loss, in contrast to the reduced sound frequency range of *Neurod1CKO* ^19^, responses for the entire measured frequency range were detected in *Isl1CKO*. The processing of high acoustic frequencies was broadly comparable between age-matched controls and *Isl1CKO*, indicating some preservation of peripheral neuronal activity. Specifically, high frequencies above 28 kHz were comparable between the control and *Isl1CKO* mice, as shown by ABR measurements and extracellular electrophysiological recordings of IC neuronal activity (Fig. 4b and 8d, e).

As a result of disorganized primary auditory neurons with derailed central projections, the characteristics of persistent auditory function in the IC were altered with worsened tuning capabilities of IC units and their increased spontaneous activity and threshold elevations and decreased dynamic range. The peripheral deficit in sound encoding results in abnormal auditory behavior of *Isl1CKO*. Although no significant differences of ABR thresholds at 32 kHz were observed between *Isl1CKO* and control mice, indicating retained hearing function, the startle reactions of *Isl1CKO* at 32 kHz were reduced. Plasticity of the startle response is also evident in the PPI responses of *Isl1CKO* mice, in which a weak prestimulus suppresses the response to a subsequent startling stimulus. *Isl1CKO* mice demonstrated PPI impairment for the pure tone of 8 kHz, reflecting abnormal sensorimotor gating due to compensatory hyperactivity of the central auditory system ^67, 81^.

Additionally, compared to control mice, DPOAE responses of *Isl1CKO* were reduced, indicating dysfunction of cochlear amplification. The OHCs of the organ of Corti play a central role in the active enhancement of sound-induced vibration. For given OHCs, amplification only occurs at a precise frequency, and thus, this mechanism provides a sharpening of the tuning curve and improves frequency selectivity ^82^. Nevertheless, DPOAE analysis showed that some function was preserved in high-frequency OHCs in the *Isl1CKO* cochlea. Frequencies above 28 kHz are located at the basal half of the mouse cochlea, from the mid-base to the basal end ^49^, which correspond to the most preserved distribution of sensory neurons in the area of the spiral ganglion of *Isl1CKO* mice (Fig. 1, 5). Usually, decreased DPOAE amplitudes indicate loss and dysfunction of OHCs ^76, 82, 83, 84^. Since *Neurod1^Cre^* is not expressed in sensory cells in the cochlea, any changes in the development of OHCs represent secondary effects of *Isl1* elimination in neurons. Accordingly, cochlear amplification deficits in *Isl1CKO* correlated with the reduced and disorganized innervation of OHCs (Fig. 2). The medial olivocochlear efferents innervate OHCs from the brainstem, representing a sound-evoked negative feedback loop suppressing OHC activity ^85^. The formation of efferent intraganglionic spiral bundle was altered in *Isl1CKO*, indicating changes in efferent innervation in the cochlear region. Besides efferents, OHCs are innervated by the type II SGNs ^86, 87^. Although the function of type II SGNs remains obscure, it is clear that these neurons are involved in auditory nociception ^88^, and may also constitute the disputed sensory drive for the olivocochlear efferent reflex ^89, 90^. As *Isl1* is expressed in both type I and type II SGNs during inner ear development ^4^, characteristics of both neuronal types might likely be affected in *Isl1CKO*.

The unexpected close-to-normal hearing function and auditory signal processing at high frequencies suggest some preservation of tonotopic organization in the *Isl1CKO* cochlea for the propagation of acoustic information. Correspondingly, our 3D images of the *Isl1CKO* cochlea show that some neurons in the base are in their proper position in the Rosenthal’s canal, forming a segment of the spiral ganglion. These unexpected results may reflect a timeline of sequential neuronal differentiation from the base to the apex, as the first differentiated neurons are in the base and the last SGNs undergo terminal mitosis in the apex ^91^. As the cochlea is extending, differentiating neurons migrate along the cochlea to form the spiral ganglion. This process is completely disrupted in the *Isl1CKO* cochlea, and many neurons are concentrated in a conical shaped central structure, the modiolus, with only a portion of neurons in the Rosenthal’s canal. Notably, the functional and spatial organization of SGNs may differ in transcription factor networks required for their differentiation programs. For example, some *Neurod1* lacking neurons survive, form a rudimental cochlear ganglion, and establish bipolar connections to their targets in *Neurod1* null mice ^11^ or the otocyst-deleted *Neurod1* conditional mutant ^17^. Consistent with these findings, neurons forming a segment of the spiral ganglion in the Rosenthal’s canal of *Isl1CKO* may not require ISL1 for their differentiation and/or the migration mode. Another possibility is that *Neurod1^Cre^* may produce a delayed *Isl1* deletion, resulting in a close to-normal spatial organization of SGNs and auditory system function at high frequencies. A different Cre driver generating an earlier recombination of *Isl1* in the otocyst might be used to address the high-frequency hearing phenotype of *Isl1CKO*.

Another secondary effect of *Isl1* elimination in neurons likely contributing to the hearing deficit of *Isl1CKO* is a shortened cochlea accompanied by the disorganized apical epithelium. Somewhat similar phenotypes of a shortened cochlea were previously reported for deletion mutants of *Neurod1* and *Neurog1*, key transcription factors for inner ear neuronal development ^11, 18, 19, 91^. A comparable effect of cochlear length reduction was observed following the loss of *Foxg1* ^92^ and *Lmx1a* ^93, 94^. Although mechanisms affecting the cochlear extension are unknown, it is clear that these confounding features of the *Isl1CKO* phenotype would consequently impact mechanical and neural tuning from the base to the apex of the cochlea and the ability to perform time-frequency processing of sound.

A limitation of our study is that we only assessed the functional role of *Isl1* in the inner ear neurons. The expression pattern of ISL1 indicates that ISL1 may control the development of multiple inner ear progenitors ^21^. Notably, the elimination of *Isl1* in developing neurons is only a part of the story. ISL1 is highly expressed in inner ear neurons throughout the development and in postnatal auditory neurons (Supplementary Figure 2) ^4, 18, 21, 22, 95^. Consistent with our results, the expression profile indicates an important role of ISL1 for the terminal differentiation of neurons, and for the function and maintenance of differentiated neurons. In contrast, the expression of ISL1 in sensory precursors is downregulated as hair cell differentiation is initiated and thus, ISL1 is not detected in the differentiated sensory hair and supporting cells in vestibular and cochlear epithelia ^21^. As the expression of ISL1 precedes the induction of ATOH1 and sensory cell differentiation, it is conceivable that ISL1 may play a role in the specification of sensory fate and the regulation of the initial sequential events in sensory precursor development. Further studies will be needed to fully uncouple regulatory mechanisms in inner ear development by targeted eliminations of *Isl1* in sensory precursors and in both neuronal and sensory precursor populations.

Altogether, our study provides compelling evidence that ISL1 is a critical regulator of SGN development, affecting neuronal migration, pathfinding abilities to form cochlear wiring, and central axonal projections. As such, ISL1 represents an essential factor in the regulation of neuronal differentiation to produce functional neurons in cell-based therapies and stem cell engineering ^96, 97^. Additionally, this unique model contributes to our understanding of how disorganization of the neuronal periphery affects information processing at higher centers of the central auditory pathway at the physiological and behavioral levels.

## Methods

### Experimental animals

All experiments involving animals were performed according to the Guide for the Care and Use of Laboratory Animals (National Research Council. Washington, DC. The National Academies Press, 1996). The design of experiments was approved by the Animal Care and Use Committee of the Institute of Molecular Genetics, Czech Academy of Sciences. The mice were housed in 12-hour light/dark cycles and were fed *ad libitum*. To generate *Isl1CKO* (the genotype *Neurod1^Cre^;Isl1^loxP/loxP^*), we cross-bred floxed *Isl1* (*Isl1^loxP/loxP^; Isl1^tm2Sev^/J*, # 028501, Jackson Laboratory)^25^ and *Neurod1^Cre^* transgenic mice (Tg(Neurod1-cre)1Able/J, # 028364, Jackson Laboratory), which were generated by pronuclear injection of the *Neurod1-cre* BAC construct that carries Cre-sequence downstream of the translational initiation codon ATG of the *Neurod1* gene ^24^. Heterozygous animals, *Neurod1^Cre^;Isl1^+/loxP^* were viable, born in appropriate Mendelian ratios, and were phenotypically indistinguishable from control (Cre negative) littermate mice. As control mice, we used mice with the genotype Cre negative, *Isl1^loxP/loxP^*, and *Isl1^+/loxP^*. The mouse line *Neurod1^Cre^* was also bred with Cre-reporter tdTomato line (*TomatoAi14*, B6.Cg-*Gt(ROSA)26Sor^tm14(CAG-tdTomato)Hze^*, # 7914 Jackson Laboratory). Using tdTomato reporter for our analyses, we compared the reporter control-*Ai14* (genotype: *Neurod1^Cre^;Isl1^+/loxP^;TomatoAi14*) and reporter *Isl1CKO-Ai14* (genotype: *Neurod1^Cre^;Isl1^loxP/loxP^;TomatoAi14*). We performed PCR genotyping on tail DNA. We used both males and females for experiments. Lines are a mixed C57BL/6/sv129 background. Phenotyping and data analyses were performed blind to the genotype of the mice.

### Morphological evaluation of the cochlea, the CN, and the IC

Dissected ears were fixed in 4% paraformaldehyde (PFA) in PBS. For vibratome sections, samples were embedded in 4% agarose and sectioned at 80 μm using a Leica VT1000S vibratome. Vibratome sections, whole inner ears, or whole embryos were defatted in 70% ethanol and then rehydrated and blocked with serum, as described previously ^20, 77^. Samples were then incubated with primary antibodies at 4°C for 72 hours. The primary antibodies used were: rabbit anti-Myosin 7a (Myo7a; Proteus BioSciences 25-6790, 1:500), mouse anti-acetylated α-tubulin (tubulin; Sigma-Aldrich T6793, 1:400), rabbit anti-calretinin (Santa Cruz Biotechnology sc-50453, 1:100), goat anti-prestin (Santa Cruz Biotechnology sc-22692, 1:50), rabbit anti-parvalbumin (Abcam ab11427, 1:2000), mouse anti-VGLUT1 (Merck MAB5502, 1:200), rabbit anti-NeuN (Abcam ab177487, 1:500), mouse anti-Isl1 (Developmental Hybridoma Bank 39.3F7 or 39.4D5, 1:130), mouse anti-C-terminal binding protein 2 (CtBP2; BD Biosciences 612044, 1:200), rabbit anti-cleaved Caspase-3 (Cell Signaling Technology 9661, 1:100), goat anti-Neurod1 (Santa Cruz Biotechnology sc-1084, 1:100), anti-β tubulin III (Tuj1; BioLegend 801202, 1:500), and rabbit anti-Sox10 (Abcam ab155279, 1:250). After several PBS washes, secondary antibodies were added and incubated at 4°C for 24 hours. The secondary antibodies Alexa Fluor^®^ 488 AffiniPure Goat Anti-Mouse IgG (Jackson ImmunoResearch Laboratories 115-545-146), Alexa Fluor^®^ 594 AffiniPure Goat Anti-Rabbit IgG (Jackson ImmunoResearch Laboratories 111-585-144), DyLight488-conjugated AffiniPure Mouse Anti-Goat IgG (Jackson ImmunoResearch Laboratories 205-485-108), Alexa Fluor^®^ 647-conjugated AffiniPure Donkey Anti-Goat IgG (Jackson ImmunoResearch Laboratories 705-605-147), Alexa Fluor^®^ 594-conjugated AffiniPure Donkey Anti-Rabbit IgG (Jackson ImmunoResearch Laboratories 711-585-152), and Alexa Fluor^®^488-conjugated AffiniPure Donkey Anti-Mouse IgG (Jackson ImmunoResearch Laboratories 715-545-150) were used in 1:400 dilution. Nuclei were stained by Hoechst 33258 (Sigma-Aldrich 861405, 1:2000). Samples were mounted in Aqua-Poly/Mount (Polysciences 18606) or prepared Antifade medium. Images were taken on Zeiss LSM 880 NLO inverted confocal microscope, Nikon CSU-W1 spinning disk confocal microscope, and Carl Zeiss AxioZoomV16 macroscope. NIS-Elements, ImageJ, and ZEN software were used for image processing.

The length of the organ of Corti was measured using the “Measure line” ImageJ plugin. The volumes of the CN and IC were established by analyzing parallel, serial, equally spaced 80 μm coronal vibratome sections through the brain (n = 5 *Isl1CKO* and n = 5 control mice). The CN and IC areas were determined in each section using ImageJ, and the volume of the organs was calculated. Volumes of organs were adjusted to the body weight. Whole-mount anti-tubulin labeling of the cochlea was used to measure the length and density of the radial fibers. The evaluation of the innervation was done separately for each region of the cochlea: the apex, mid-apex, mid-base, and base. Due to the disorganization of innervation in the apex, we only evaluated the cochlear base, mid-base, and mid-apex fiber density. The density of the radial fibers was expressed as the percentage of a positive area in the measured area of 152 x 66 μm^2^ using the “Threshold” function of ImageJ. The length of the radial fibers was measured from the whole-mount anti-tubulin and Myo7a immunolabeled cochlea confocal images. For each genotype, we measured 3 samples, and in all four regions of the cochlea (the apex, mid-apex, mid-base, and base), we measured the length of the fibers in 3 radial fiber bundles from the IHCs to the IGSB. To compare how many neurons are correctly located in the Rosenthal’s canal in *Isl1CKO*, we used NeuN immunolabeled whole-mount cochlea (n = 4 per genotype). Cells were manually counted in 40x magnification images in 5 Z-stacks using the “Cell counter” function of ImageJ; the area of the Rosenthal’s canal with tdTomato^+^ neurons was measured using “Threshold”, “Polygon selection” and “Measure” functions of ImageJ to calculate the relative number of neurons for the control and *Isl1CKO* per Rosenthal’s canal volume. For ribbon evaluations, IHCs were counted based on the calretinin-stained cell bodies and anti-CtBP2 labeled synapses. CtBP2^+^ presynaptic ribbon structures were manually counted in 10 individual IHCs in each cochlear region: the apex, mid-apex, mid-base, and base. A total of 6 mutants and 6 control cochleae were evaluated except for 1 apex in the control group, which was lost during immunolabeling. The apoptosis of neurons was quantified using immunohistochemistry with anti-cleaved Caspase-3. Caspase-3 positive cells were manually counted in the inner ear ganglia labeled by the combination of anti-Cre and anti-Neurod1 in the whole-mounted head at 60x magnification at E10.5. For E12.5, Caspase-3^+^/tdTomato^+^ neurons were manually counted in the whole-mounted inner ear at 60x magnification. The area of E12.5 inner ear ganglia was measured using the “Threshold and polygon selection” function of ImageJ. For E14.5, the cochlea was sectioned, and Caspase-3^+/tdTomato+^ neurons were manually counted in all sections at 60x magnification. Three embryos per genotype were used for each embryonic day. All apoptotic cell counting was done using the “Cell counter” function of ImageJ. The density of spherical bushy cells in the VCN, 3 control and 3 *ISL1CKO* 2-month-old mice were used. Bushy cells were labeled by anti-parvalbumin in 80 μm vibratome sections, and manually counted in 2-4 sections/animal in 134.95 x 134.95 μm (63x objective). The size of vestibular sensory epithelia was measured in anti-Myo7a labeled wholemount preparations at P0. Myo7a^+^ areas were measured in 5 control and 5 *Isl1CKO* mice using “Measure” ImageJ tool. The dorsal root ganglia (DRG) size was evaluated in the sagittal sections of E14.5 embryos with expression of tdTomato reporter. The area of DRG was quantified in 6 embryos per genotype and 2 sections per embryo using the “Threshold and polygon selection” function of ImageJ.

### Light-sheet fluorescent microscopy (LFSM) and analysis of images

The cochleae were microdissected from 2 control-*Ai14* and 2 *Isl1CKO-Ai14* mice (postnatal day P0). We used an advanced CUBIC protocol^98^ for tissue clearing to enable efficient imaging by light-sheet microscopy. Briefly, the microdissected cochlea was fixed in 4% PFA for 1 hrs, washed with PBS, and incubated in a clearing solution Cubic 1 for 7 days at 37°C. Before immunolabeling, samples were washed in PBT (0.5% Triton-X in PBS) 4 x for 30 min. In addition to *tdTomato* expression, cochlear samples were either immunolabeled using anti-NeuN (a nuclear marker of differentiated neurons) or anti-Myo7a (a hair cell marker) antibodies. Samples were stored before imaging in Cubic 2 at room temperature. Zeiss Lightsheet Z.1 microscope with illumination objective Lightsheet Z.1 5x/0.1 and detection objective Dry objective Lightsheet Z.1 5x/0.16 was used for imaging at the Light Microscopy Core Facility of the Institute of Molecular Genetics of the Czech Academy of Sciences. IMARIS software v8.1.1 (Bitplane AG, CA, USA) was used for image processing.

### Isolation of genetically labeled neurons and library construction

Spiral ganglia were micro-dissected in Dulbecco’s PBS on ice from E14.5 embryos of either sex from four litters; each sample contains spiral ganglia from both inner ears of the individual embryo. Spiral ganglia were incubated in 300 μl of lysis solution (0.05% trypsin, 0.53mM EDTA in Dulbecco’s PBS) in 37 °C, shaking at 900 RPM for 5 min. The lysis was stopped by adding 600 μl of FACS buffer (10mM EGTA, and 2% FBS in Dulbecco’s PBS). After spinning down the samples at 800 G, 4 °C for 10 min, the supernatant was removed, and cell pellets were resuspended in 500 μl of ice-cold FACS buffer. Immediately before sorting, cells were passed through a 50 μm cell sieve (CellTrics™, Sysmex Amercica Inc.) into a sterile 5 ml polystyrene round-bottom falcon to remove clusters of cells and kept on ice. TdTomato^+^ neurons were sorted using a flow cytometer (BD FACSAria™ Fusion), through a 100 μm nozzle in 20 psi, operated with BD FACSDiva™ Software (Supplementary Fig. 9). 100 sorted cells were collected into individual wells of 96-well plate containing 5 μl of lysis buffer of NEB Next single-cell low input RNA library prep kit for Illumina (New England Biolabs #E6420). Plates were frozen immediately on dry ice and stored at −80 °C. The total time from euthanasia to cell collection was ~3 hrs.

RNAseq-libraries were prepared from 6 samples per each genotype, reporter control-*Ai14* and *Isl1CKO-Ai14* mutant, and each sample contained 100 tdTomato^+^ neurons. Following the manufacturer’s instructions, the NEB Next single-cell low input RNA library prep kit for Illumina was used for cDNA synthesis, amplification, and library generation ^99^ at the Gene Core Facility (Institute of Biotechnology CAS, Czechia). Fragment Analyzer assessed the quality of cDNA libraries. The libraries were sequenced on an Illumina NextSeq 500 next-generation sequencer. NextSeq 500/550 High Output kit 75 cycles (Illumina #200024906) were processed at the Genomics and Bioinformatics Core Facility (Institute of Molecular Genetics CAS, Czechia).

### Computational analysis of RNAseq data

RNA-Seq reads in FASTQ files were mapped to the mouse genome using STAR [version 2.7.0c ^100^] GRCm38 primary assembly and annotation version M8. The raw data of RNA sequencing were processed with a standard pipeline. Using cutadapt v1.18^101^, the number of reads (minimum, 32 million; maximum, 73 million) was trimmed by Illumina sequencing adaptor and of bases with reading quality lower than 20, subsequently reads shorter than 20 bp were filtered out. TrimmomaticPE version 0.36 ^102^. Ribosomal RNA and reads mapping to UniVec database were filtered out using bowtie v1.2.2. with parameters -S -n 1 and SortMeRNA ^103^. A count table was generated by Rsubread v2.0.1 package using default parameters without counting multi mapping reads. The raw RNAseq data were deposited at GEO: # GSE182575 study (https://www.ncbi.nlm.nih.gov/geo/query/acc.cgi?acc=GSE182575). DESeq2 [v1.26.0 ^104^] default parameters were used to normalize data and compare the different groups. Differentially expressed genes were identified based on an adjusted P-value p_adj_ < 0.05, FC > 2, and a base mean ≥ 50 was applied to identify differentially expressed genes between *Isl1CKO* mutant and control neurons. The functional annotation of the differentially expressed genes was performed using GOTermFinder ^105^.

#### Enrichment mapping

The enrichment of the functional categories and functional annotation clustering of the differentially expressed genes was performed using g: Profiler ^106^ input using version e104_eg51_p15_3922dba with g: SCS multiple testing correction methods applying a significance threshold of 0.05. In contrast, no electronic GO annotations were used. Only Biological Processes (BP) data underwent further processing. Complete query details are available in Query info tabs in Supplementary Data 2. The resulting GEM and combined GMT files were loaded into Cytoscape ^107^ plugin “EnrichmentMap” ^108^ using 0.01 FDR q-value cutoff to generate a network. Edge cutoff was set to 0.6, and nodes were filtered by gs_size<1800. Five GO terms forming solitary nodes, or a pair of nodes, were excluded (listed in Supplementary Data 2). Further adjustments were made in yFiles Layout Algorithms, Legend Creator (Cytoscape plugins), and Inkscape (Inkscape Project, 2020).

### Quantitative real-time PCR

Total RNA was isolated from both inner ears of the embryo at E14.5 using TRI Reagent (Sigma-Aldrich T9424). We used 8 embryos for *Isl1CKO* and 7 embryos for the control group from three litters. RNA from both inner ears of one embryo represented one sample. RNA samples (0.5 μg of total RNA) were processed and analyzed as previously described ^18, 77^. Briefly, following RT, quantitative qPCR was performed with initial activation at 95 °C for 120 s, followed by 40 cycles at 95 °C for 15 s, 60 °C for 30 s, and 72 °C for 30 s using the CFX384™ Real-Time PCR Detection System (Bio-Rad Laboratories). The primer sequences (*Lhx1, Lhx2, Cdh7, Nhlh2, Ntrk2, Ntrk3, Ntng1*) were obtained from (pga.mgh.harvard.edu/primerbank/) or designed using Primer3 tool (*Dcc, Epha5, Gata3, Prdm8, Robo2, Slit2, Tbx3, Uncx*). The primer sequences are listed in Supplementary Table 2. Relative mRNA expression was calculated using the –ΔΔCq method with *Hprt1* as a reference gene. GraphPad Prism software was used for statistical analysis.

### Lipophilic Dye Tracing

We studied the innervation pattern in whole or dissected ears using lipophilic dye tracing in aldehyde-fixed tissues as previously described ^109^. At least three mutants and similar numbers of control littermates of both sexes were used for each evaluation. Filter strips loaded with NeuroVue colored lipophilic dyes were inserted into the cochlear apex, base, and vestibular end-organ utricle. After allowing appropriate time for diffusion of the lipophilic tracer (between 48-120 hours), we prepared the ears as whole mounts in glycerol on a glass slide, using appropriate spacers to avoid distortion, and imaged them using a Leica SP8 confocal microscope. Images were compiled into plates to show the most pertinent details using Corel Draw. Only general image modifications such as contrast, or brightness adjustments were used to enhance the visual appeal without affecting the scientific content.

### Hearing function evaluation

Auditory brainstem response (ABR) and distortion product otoacoustic emissions (DPOAEs) tests were carried out on mice under general anesthesia with 35 mg/kg ketamine (Calypso 50 mg/ml) and 6 mg/kg xylazine (Xylapan 20 mg/ml) in saline to give an application volume of 7 ml/kg body weight via subcutaneous injection, maintained on a temperature-regulated blanket in a soundproof room.

#### Distortion product otoacoustic emissions

For DPOAE recording were tested *Isl1CKO* (n = 10) and control mice (n = 14). Cubic (2 F1–F2) distortion product otoacoustic emissions over an F2 frequency range from 4 to 39 kHz were recorded with a low-noise microphone system (Etymotic probe ER-10B+, Etymotic Research). Acoustic stimuli (ratio F2/F1 = 1.21, F1 and F2 primary tone levels of L1/L2 = 70/60 dB) were presented to the ear canal with two custom-made piezoelectric stimulators connected to the probe with 10-cm-long silastic tubes. The signal from the microphone was analyzed by the TDT System III (RP2 processor, sampling rate 100 kHz) using custom-made MATLAB software. DPOAEs were recorded in the animals’ ears at individual frequencies over the frequency range 4–39 kHz with a resolution of ten points per octave. All the experiments and analyses were done with no information on the genotype.

#### Auditory brainstem response

For auditory brainstem response (ABR) recording (n = 10 *Isl1CKO* and n = 11 control mice), an active electrode was placed subcutaneously on the vertex and ground and reference electrodes in the neck muscles. Responses to tone bursts (3 ms duration, 1 ms rise/fall times, frequencies of 2, 4, 8, 16, 32, and 40 kHz) and clicks of different intensity were recorded. Acoustic stimuli were conveyed to the animal in free-field conditions via a two-way loudspeaker system (Selenium 6W4P woofer and RAAL70-20 tweeter) placed 70 cm in front of the animal’s head. The signal was processed with a TDT System III Pentusa Base Station and analyzed using BioSig™ software. The response threshold to each frequency was determined as the minimal tone intensity that still evoked a noticeable potential peak in the expected time window of the recorded signal. The amplitude and latency of ABR peaks I-V were determined using BioSig software (Tucker Davis Technologies). Central compensation of neuronal responsiveness (central gain) was calculated using ABR wave IV to I amplitudes.

### Extracellular recording of the neuronal activity in the IC

For extracellular recording in the IC, we evaluated *Isl1CKO* (n = 12, 624 units) and control mice (n = 8, 395 units). The surgery and extracellular recording in the IC were performed in mice anesthetized with 35 mg/kg ketamine (Calypsol 50 mg/ml) and 6 mg/kg xylazine (Xylapan 20 mg/ml) in saline via subcutaneous injection. Approximately every hour, supplement subcutaneous injections of one-half of the original dose of the anesthetics were administered to keep a sufficient level of anesthesia, judged by a positive pedal and palpebral (toe-pinch) reflex and movement of the whiskers. Respiratory rate, and heart rate, were monitored. An incision was made through the skull’s skin to access the IC, and underlying muscles were retracted to expose the dorsal skull. A holder was glued to the skull, and small holes were drilled over both ICs. Neuronal activity (multiple units) in the IC was recorded using a 16-channel, single shank probe (NeuroNexus Technologies) with 50 or 100 μm between the electrode spots. The obtained signal from the electrode was amplified 10000 times, band-pass filtered over the range of 300 Hz to 10 kHz, and processed by a TDT System III (Tucker Davis Technologies) using an RX5-2 Pentusa Base Station. Individual spikes from the recorded signal were isolated online based on amplitude discrimination and analyzed with BrainWare software (v. 8.12, Jan Schnupp, Oxford University). Subsequent discrimination of spikes from the recorded data and their sorting according to the amplitudes of the first positive and negative peaks were performed off-line and was used to sort action potentials (spikes) among single units. The recorded data were processed and analyzed using custom software based on MATLAB. The stimulation signals were generated using a TDT System III with the RP 2.1 Enhanced Real-Time Processor. Acoustic stimuli were delivered in free-field conditions via a two-driver loudspeaker system (Selenium 6W4P woofer and RAAL70-20 tweeter) placed 70 cm in front of the animal’s head.

#### Frequency-intensity mapping

To determine the neuronal receptive fields, pure tones (frequency 2 - 40 kHz with 1/8 octave step, 60 ms duration, 5 ms rise/fall times, various intensity with 5dB step) were presented in a random order, each stimulus appearing three times. A discrete matrix corresponding to the response magnitude evoked by each of the frequency-intensity combinations was thereby obtained, smoothed using cubic spline interpolation, and used for extraction of the basic parameters: the excitatory response threshold (the lowest stimulus intensity that excited the neuron, measured in dB SPL), the characteristic frequency (CF) - the frequency with the minimal response threshold, measured in Hz, and the bandwidth of the excitatory area 20 dB above the excitatory threshold, expressed by quality factor Q (Q = CF/bandwidth).

#### Rate intensity function of the IC neurons

Neuronal responses to broadband noise (BBN) bursts of variable intensity (10 dB steps, 50 repetitions) were used to construct the rate intensity function (RIF). A 100% scale was assigned to the neuron’s total range of response amplitudes, 0% corresponding to spontaneous activity and 100% corresponding to its maximum response magnitude ^110^. The two points of interest are R10 and R90, which correspond to 10 and 90% of this scale, respectively. R10, describing the starting point of the RIF’s rise, was taken as the BBN response threshold. RIFs were further used to evaluate the following parameters: the dynamic range (DR) of the RIF: DR = S90–S10); and the maximum response magnitude. Spontaneous activity of the IC neurons was determined at the 0dB SPL BBN stimulation.

#### Temporal properties of the IC neurons

We used trains of five clicks at an intensity of 70 dB SPL for control and 80 dB SPL for *Isl1CKO* mice with various inter-click intervals (100, 50, 30, 20, and 15 ms). We calculated the vector strength (VS) values and the Rayleigh statistics for each spike pattern; only responses with at least 5.991 were significantly considered phase-locking (Zhou and Merzenich, 2008). The VS quantifies how well the individual spikes are synchronized (phase-locked) with a periodic signal.

### Behavioral tests of motor coordination and balance

All testing was carried out during the light cycle and after a minimum 30-min acclimatization. *Air-righting reflex*. The vestibular function was evaluated by the ability of the mice to right themselves in the air when held supine and dropped onto a soft surface from a height of 50 cm^77^.

#### Rotarod

Using the rotarod apparatus (Rota Rod 47600, Ugo Basile), the time (latency) to fall off the rod rotating under continuous acceleration was recorded. During the acclimatization period, mice with their heads in the direction of rotation were loaded on the rotarod at an initial speed of 4 rpm. This speed was maintained for 2 min and, if mice fell during this period, they were placed back on the rotarod. The drum was slowly accelerated from 4 to 40 rpm for a maximum of 300 s for each trial for the experimental measurements. The latency to fall off the rotarod within 300 s was recorded. If the mouse clung to the grip of the rotating rod and failed to resume normal performance for three consecutive revolutions, the sensor was manually triggered. Mice were tested in three consecutive trials in one session per day with a 15-min rest period between each trial.

### Behavioral acoustic tests

*Isl1CKO* (n = 10) and control (n = 7) mice were used at 2-3 months of age. All behavioral tests were performed in a sound-attenuated chamber (Coulbourn Habitest, model E10-21) located in a soundproof room. Each mouse was placed in a wire mesh cage on a motion-sensitive platform inside the box during the testing. The mouse’s reflex movements to sound stimuli were detected and transformed to a voltage signal by the load-cell response sensing platform. An amplified voltage signal was acquired and processed using a TDT system 3 with a Real-Time Processor RP 2 (Tucker Davis Technologies, Alachua, Fl) and custom-made software in a Matlab environment. The startle responses were evaluated in 100 ms windows, beginning at the onset of the startling stimulus. The magnitude of the response was measured as the maximal peak-to-peak amplitude of transient voltage occurring in the response window. Acoustic stimuli were generated by the TDT system (Real-Time Processor RP 2), amplified and presented via a loudspeaker (SEAS, 29AF/W), and placed inside the chamber above the animal. Stimulus presentation and data acquisition were controlled by a custom-made application in a Matlab environment. Calibration of the apparatus was performed for frequencies between 4 kHz and 32 kHz by a 1/4-inch Brüel & Kjaer 4939 microphone connected to a Brüel & Kjaer ZC 0020 preamplifier and a B&K 2231 sound level meter. During the calibration, the calibrating microphone was positioned at the animal’s head in the test cage.

*Acoustic startle reflex (ASR)* (a transient motor response to an intense, unexpected stimulus) was used to indicate the behavioral responsiveness to sound stimuli. The ASRs to 8, 16, and 32 kHz tone pips and BBN bursts (50 ms duration, 3 ms rise/fall times, varying intensity levels) were recorded. Each test session contained: a baseline trial (–10 dB SPL stimulus intensity) and 13 startle stimuli of different intensities (50, 55, 60, 65, 70, 75, 80, 85, 90, 100, 110, 115, and 120-dB SPL). The inter-trial interval varied from 15 to 50 s.

In *the prepulse inhibition (PPI)* procedure, 3 different trial types were used: a baseline trial without any stimulus, an acoustic startle pulse alone (white noise at 110 dB SPL, 50 ms, 3 ms rise/fall times), and a combination of the prepulse and startle pulse. The inter-stimulus interval between the prepulse and the startle stimulus was set to 50 ms; each trial type was presented three times. The inter-trial gap was randomized and varied from 15 to 50 s. The efficacy of the PPI of ASR was expressed as an ASR ratio in percentage, e.g., 100% corresponds to the amplitude of ASR without prepulse; smaller values of ASR ratio indicate stronger PPI. As a prepulse, we used either BBN bursts or tone pips (50 ms duration, 3 ms rise/fall time) at frequencies of 8 and 32 kHz at increasing intensities. It is expected that in the presence of prepulse, the amplitude of the following startle response decreases.

### Experimental design and statistical analyses

All comparisons were made between animals with the same genetic background, typically littermates, and we used male and female mice. The number of samples (n) for each comparison can be found in the individual method descriptions and are given in the corresponding figures. Phenotyping and data analysis was performed blind to the genotype of the mice. All values are presented either as the mean ± standard deviation (SD) or standard error of the mean (SEM). For statistical analysis, GraphPad Prism software was used. To assess differences in the mean, one-way or two-way ANOVA with Bonferroni’s multiple comparison test, multiple *t*-tests with Holm-Sidak comparison method, repeated measures ANOVA, multiple *t*-tests, and unpaired two-tailed *t*-tests were employed. Significance was determined as P < 0.05 (*), P < 0.01 (**), P < 0.001 (***) or P < 0.0001 (****). Complete results of the statistical analyses are included in the figure legends.

## Ethics approval and consent to participate

Experiments were carried out following the animal welfare guidelines 2010/63/EC of the European Communities Council Directive, agreeing with the Guide for the Care and Use of Laboratory Animals (National Research Council. Washington, DC. The National Academies Press, 1996). The design of experiments was approved by the Animal Care and Use Committee of the Institute of Molecular Genetics, Czech Academy of Sciences (protocol # 104/2019).

## Consent for publication

Not applicable.

## Data availability

All data generated or analyzed during this study are included in this published article, and Supplementary information. RNA-seq data are deposited in the publicly available Gene Expression Omnibus (GEO) repository (# GSE182575 study). Source data are provided with this paper.

## Authors’ contributions

GP, JS, BF designed and supervised the experiments. IF, KP, RB, MT, SV, MD, and BF performed experiments and analyzed the data. JC, PH, and VF prepared samples and generated 3D images. OS, SB, and LV carried RNAseq analyses. IF and KP prepared the first draft of the manuscript. GP and BF wrote the manuscript, ENY and JS reviewed the manuscript. All authors read and approved the final manuscript.

## Acknowledgments

A. Pavlinek (King’s College London) for editing the MS. We thank M. Anderova for providing TomatoAi14 reporter line. We acknowledge Imaging Methods Core Facility at BIOCEV supported by the MEYS CR (Large RI Project LM2018129 Czech-BioImaging) and ERDF (project No. CZ.02.1.01/0.0/0.0/18_046/0016045) for its support with obtaining imaging data and FACS experiments presented in this paper; Biocev GeneCore Facility for its support with gene expression/transcriptome analyses, and Biocev Animal facility (LM 2018126 Czech Centre for Phenogenomics by MEYS OP RDE CZ.02.1.01/0.0/0.0/18_046/0015861 CCP Infrastructure Upgrade II by MEYS and ESIF).

