## Supplementary Figures for "ISL1 is necessary for auditory neuron development and contributes towards tonotopic organization"

### Table of contents

1. Supplementary Figures 1-9
2. Supplementary Tables 1, 2

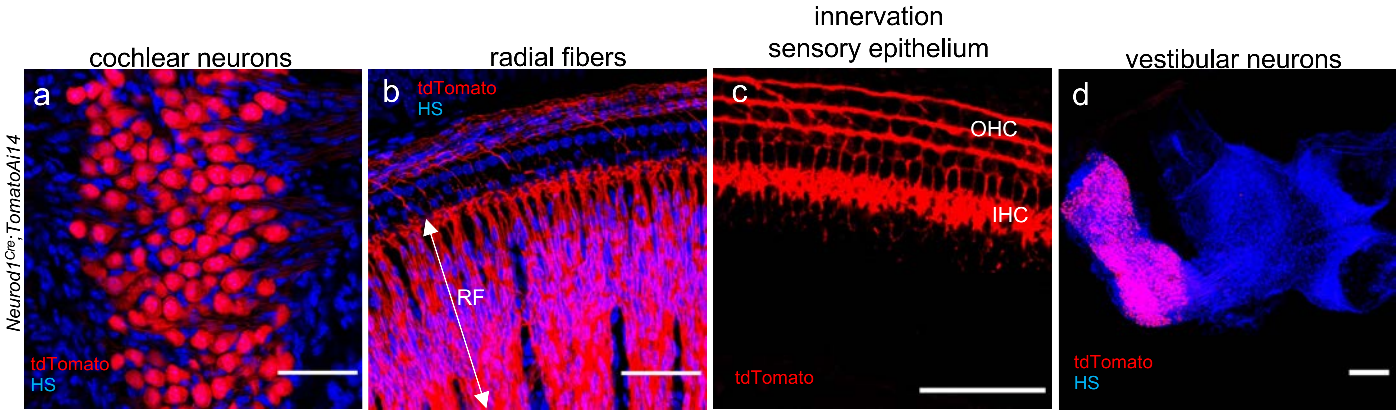


**Supplementary Fig. 1. The expression pattern of *Neurod1$^{Cre}$* shown by tdTomato reporter in the cochlea and vestibular ganglion of the *Neurod1-Cre;Ai14 tdTomato* mouse.** The confocal images from cochlear whole-mount preparations show in detail **(a)** tdTomato expression in neurons, **(b)** tdTomato$^{+}$ labeled neurites extending from neurons as radial fibers (RF) to the sensory cells, and **(c)** tdTomato$^{+}$ neurites formed three rows of outer spiral bundles beneath the outer hair cells (OHC) and the inner spiral plexus associated with the inner hair cells (IHC). **(d)** tdTomato$^{+}$ neurons of vestibular ganglion are shown in the whole-mount of the inner ear. HS, Hoechst nuclear staining. Scale bars: 50 µm (a-c), 200 µm (d).

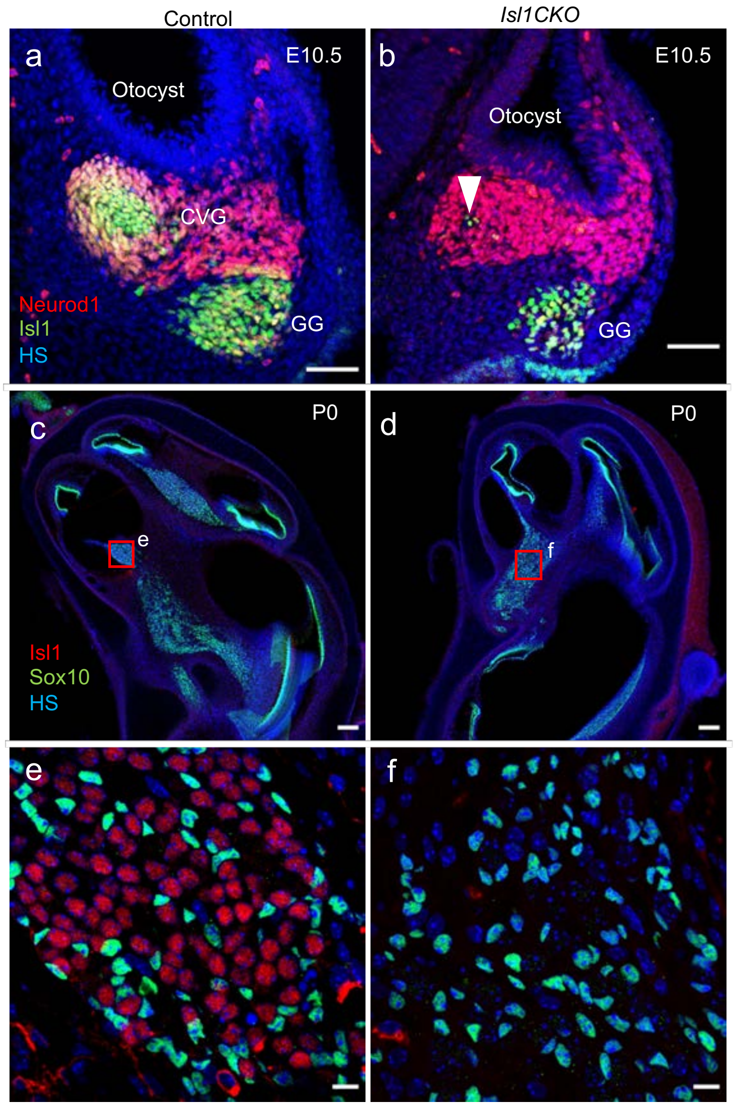


**Supplementary Fig. 2. Elimination of ISL1 shown in inner ear neurons of *Isl1CKO*. (a, b)** In E10.5 control embryos, ISL1 is expressed in differentiating neurons of cochleovestibular ganglion (CVG) and geniculate ganglion (GG), whereas ISL1 is detected only in a few neurons in the CVG of *Isl1CKO* (arrowhead). **(c-f)** At P0, ISL1 is eliminated in spiral ganglion neurons of *Isl1CKO*. Note presence of glial cells in the spiral ganglion (Sox10 is a marker of glial cells). HS, Hoechst nuclear staining. Scale bars: 50 µm (a, b); 100 µm (c, d); 10 µm (e, f).

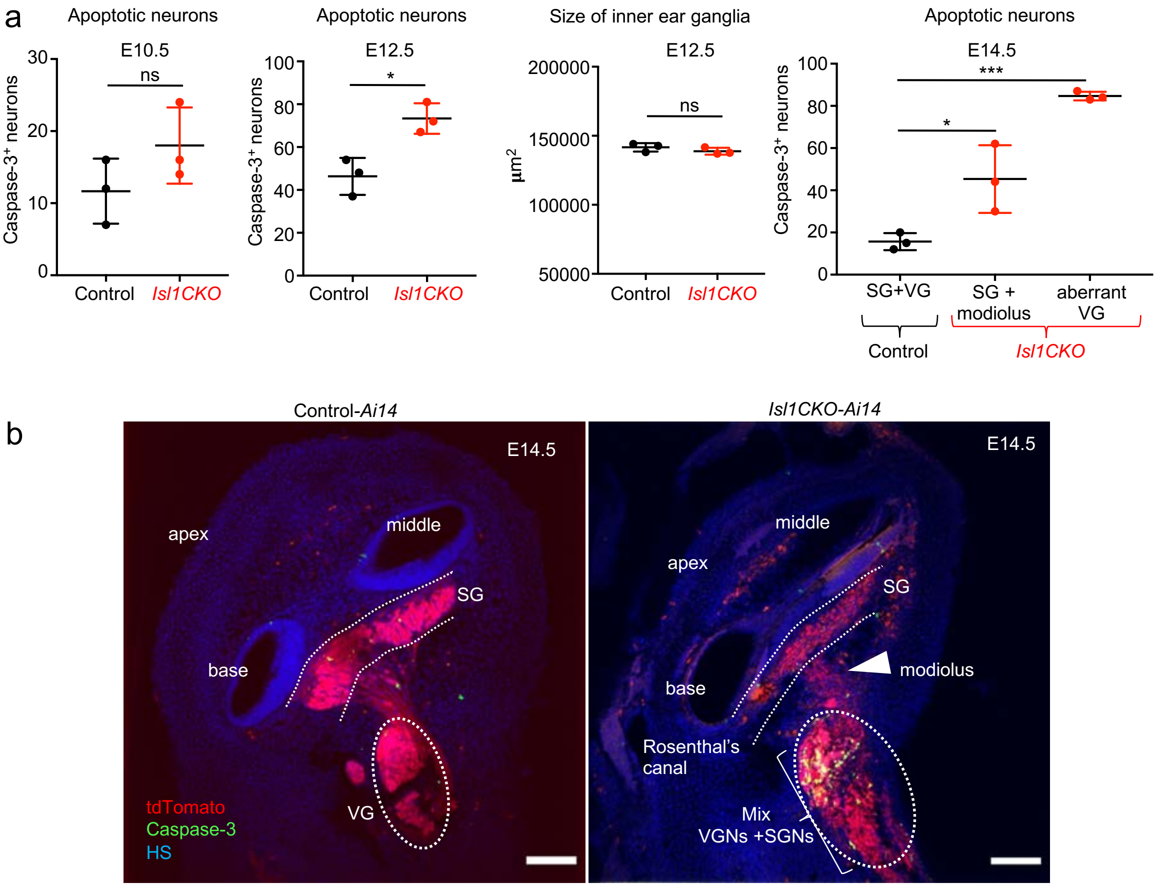


**Supplementary Fig. 3. Quantification of apoptotic neurons in the developing inner ear. (a)** The number of apoptotic neurons was manually counted in inner ear ganglia at E10.5, E12.5 and E14.5 using immunolabeling with anti-cleaved Caspase-3. The size of E12.5 inner ear ganglia is comparable between control and mutant. At E14.5, the total number of apoptotic neurons were counted in the spiral and vestibular ganglia of control embryos. In *Isl1CKO,* apoptotic neurons were counted in all areas containing neurons: the Rosenthal's canal (spiral ganglion), modiolus, and the area outside of the cochlea corresponding to the region of the vestibular ganglion in controls. Error bars represent mean ± SD (n = 3 embryos per genotype); unpaired *t*-test (E10.5, E12.5), *P = 0.0158; ns, not significant; and one-way ANOVA followed by Dunnett's multiple comparisons test (E14.5), *P = 0.0284, ***P = 0.0002. **(b)** The representative sections of the inner ear of E14.5 embryos show tdTomato$^{+}$ neurons and immunolabeled apoptotic cells by anti-cleaved Caspase-3. HS, Hoechst nuclear staining; SG, spiral ganglion; SGNs, spiral ganglion neurons; VGNs, vestibular ganglion neurons; VG, vestibular ganglion. Scale bars: 100 µm.

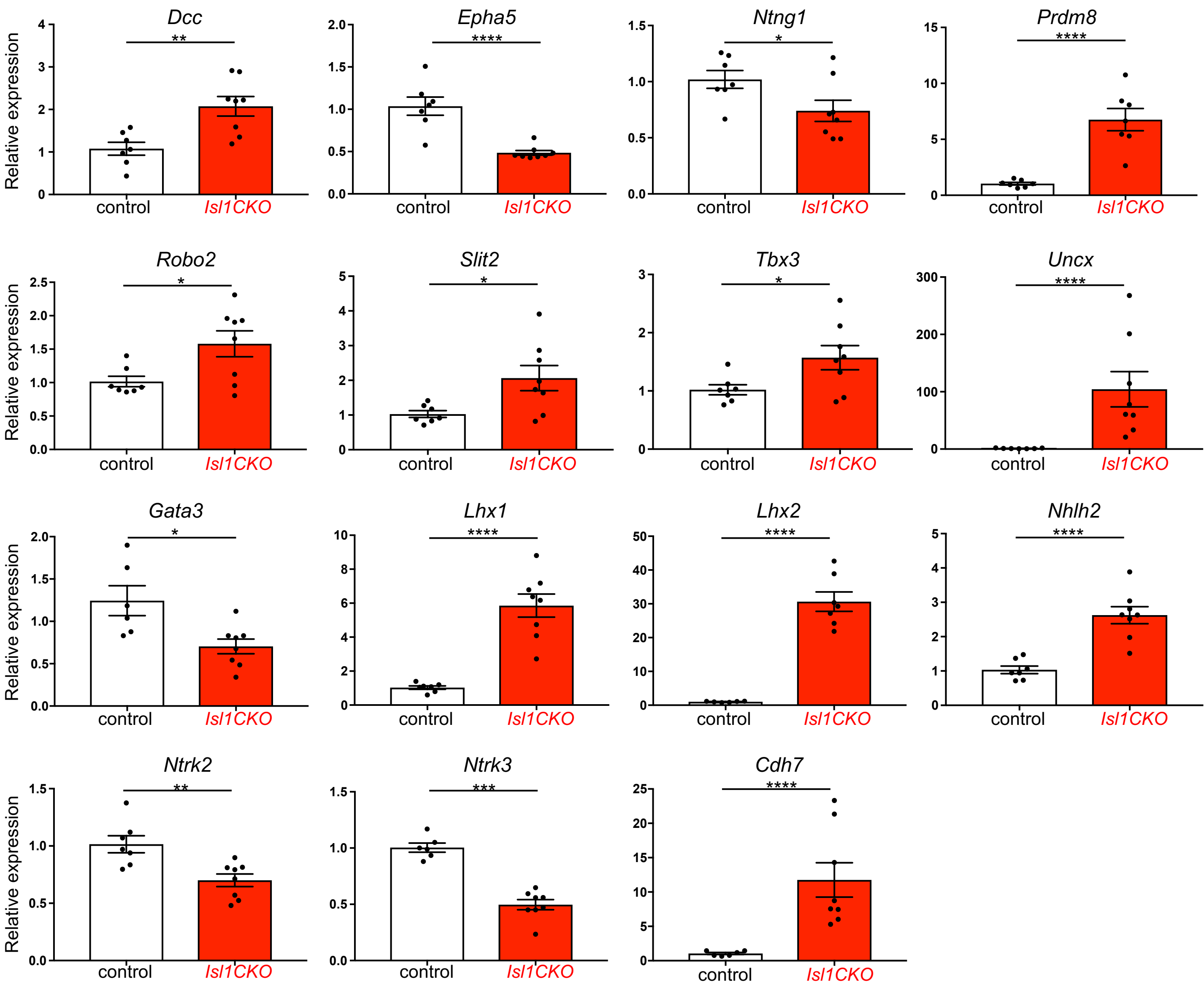


**Supplementary Fig.4. Validation of relative mRNA expression levels of selected genes by RT-qPCR.** RNA was isolated from the embryonic whole inner ear at E14.5. Data are normalized to *Hprt1* mRNA of the control gene. Data are expressed as mean ± SEM; unpaired *t*-test, *P < 0.05, **P < 0.01, ***P < 0.001, ****P < 0.0001. *Dcc*, deleted in colorectal carcinoma; *Epha5*, Eph receptor A5; *Ntng1*, netrin G1; *Prdm8*, PR domain containing 8; *Robo2*, roundabout guidance receptor 2; *Slit2*, slit guidance ligand 2; *Tbx3*, T-box 3; *Uncx*, UNC homeobox; *Gata3*, GATA binding protein 3; *Lhx1*, LIM homeobox protein 1; *Lhx2*, LIM homeobox protein 2; *Nhlh2*, nescient helix loop helix 2; *Ntrk2* and *Ntrk3*, neurotrophic tyrosine kinase receptors type 2 and type 3; and *Cdh7*, cadherin 7.

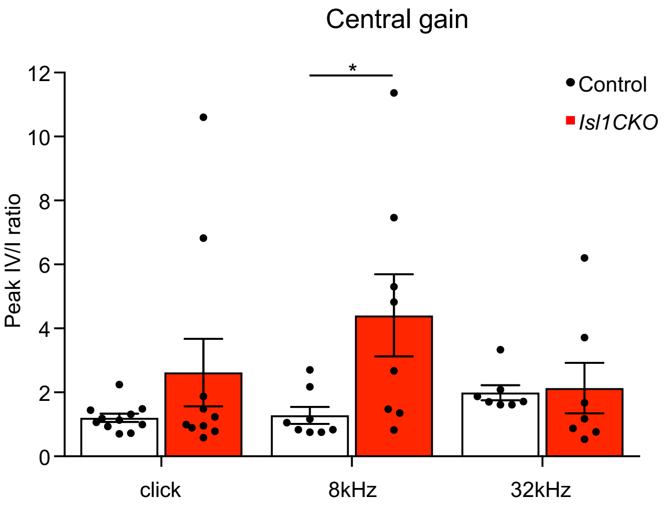


**Supplementary Fig. 5. Average ratios of amplitudes of ABR peak IV relative to peak I.** Ratios are calculated for responses to clicks at stimulus levels of 80 dB SPL and tones of frequencies 8 and 32 kHz at stimulus levels of 90 dB SPL. Data are expressed as mean ± SEM. Two-way ANOVA with Bonferroni post hoc test, *P < 0.05.

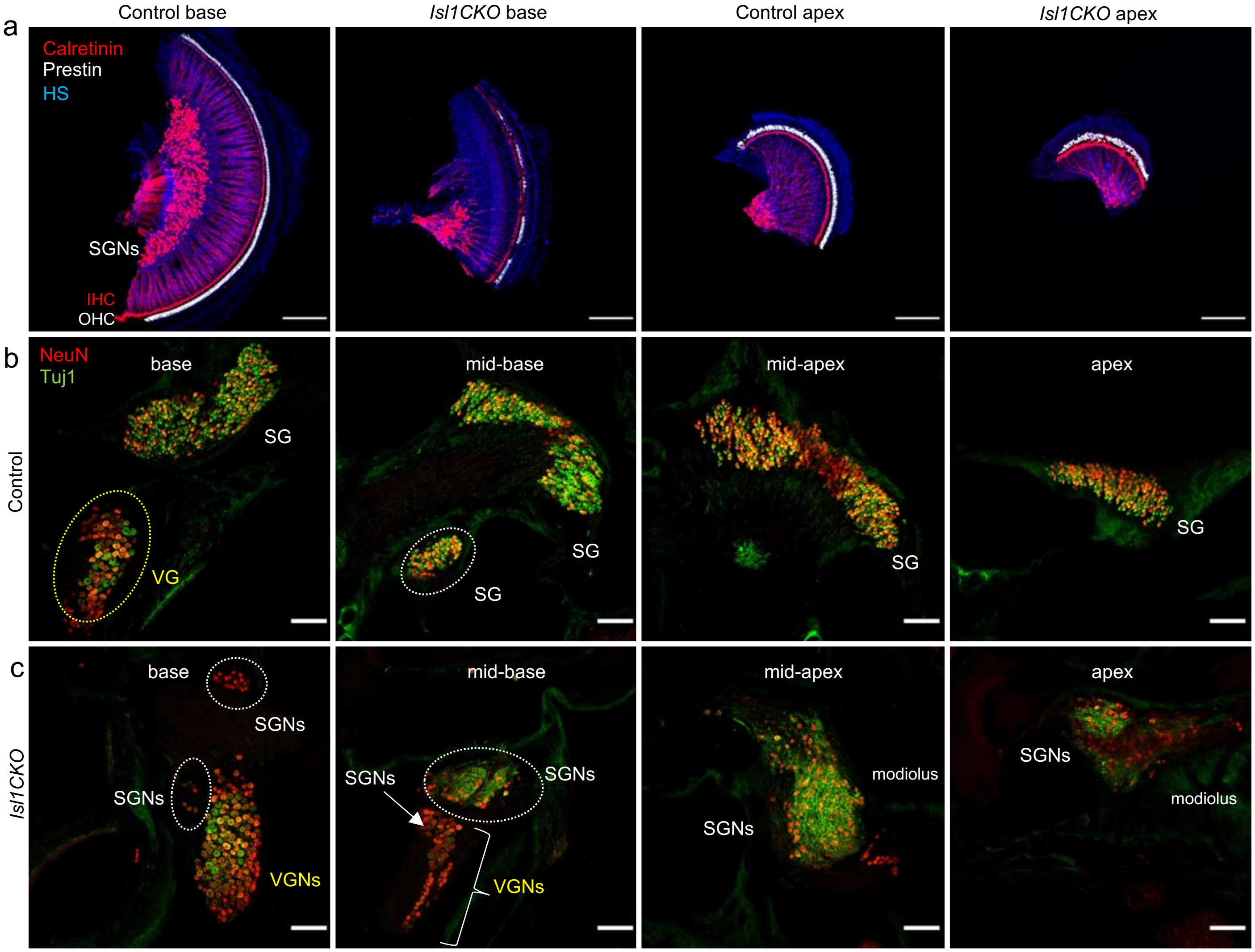


**Supplementary Fig. 6. Adult neuronal phenotype of the inner ear. (a)** Anti-calretinin labeled inner hair cells (IHC) and SGNs type Ia are shown in cochlear whole mounts of 2-month-old mice. Anti-prestin labels outer hair cells (OHC) in the sensory epithelium. **(b)** Representative images of the control inner ear sections show the location of neurons in the SG in the Rosenthal's canal and in the VG of 2-month-old-mice. Four cochlear regions, the apex, mid-apex, mid-base and base, are shown. **(c)** SGNs are misplaced in the modiolus and mixed with VGNs, based on their soma size in the adult *Isl1CKO* inner ear. HS, Hoechst nuclear staining; SG, spiral ganglion; SGNs, spiral ganglion neurons; VGNs, vestibular ganglion neurons; VG, vestibular ganglion. Scale bars: 200 μm (a), 100 μm (b, c).

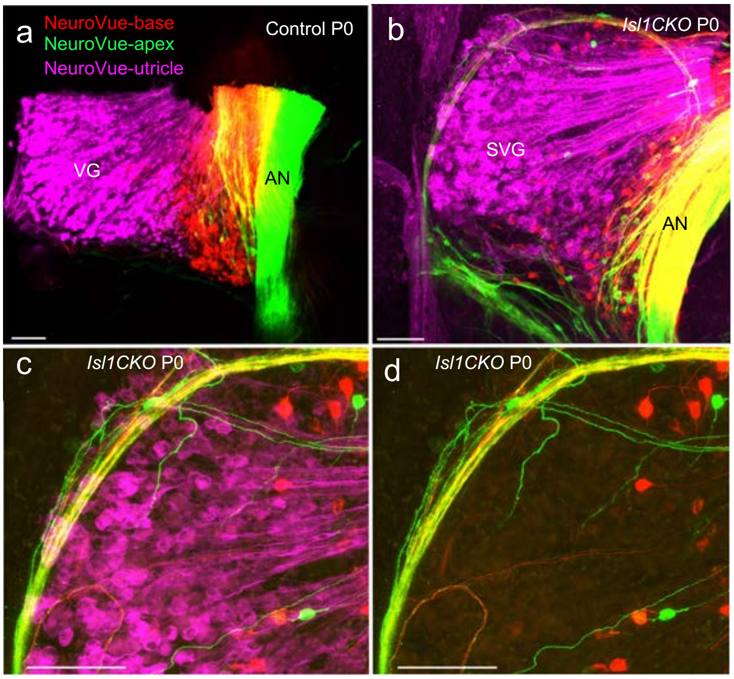


**Supplementary Fig. 7. Disorganized and unsegregated axons together with mispositioned cochlear neurons in *Isl1CKO.*** **(a)** In the P0 control, the insertions of differently colored NeuroVue lipophilic dyes into the utricle (magenta), base (red), and apex (green) labeled segregated axons in the auditory nerve (AN), and vestibular ganglion (VG) neurons and their central projections. **(b)** The same dye applications demonstrates overlayed fibers in the AN, and mixed vestibular and cochlear neurons in the spiro-vestibular ganglion (SVG) in *Isl1CKO*. **(c, d)** Higher magnification images show mislocated neurons among vestibular neurons labeled by the dye insertions into the cochlear base and apex. Note an irregular fiber loop around the SVG. Scale bars: 100 µm.

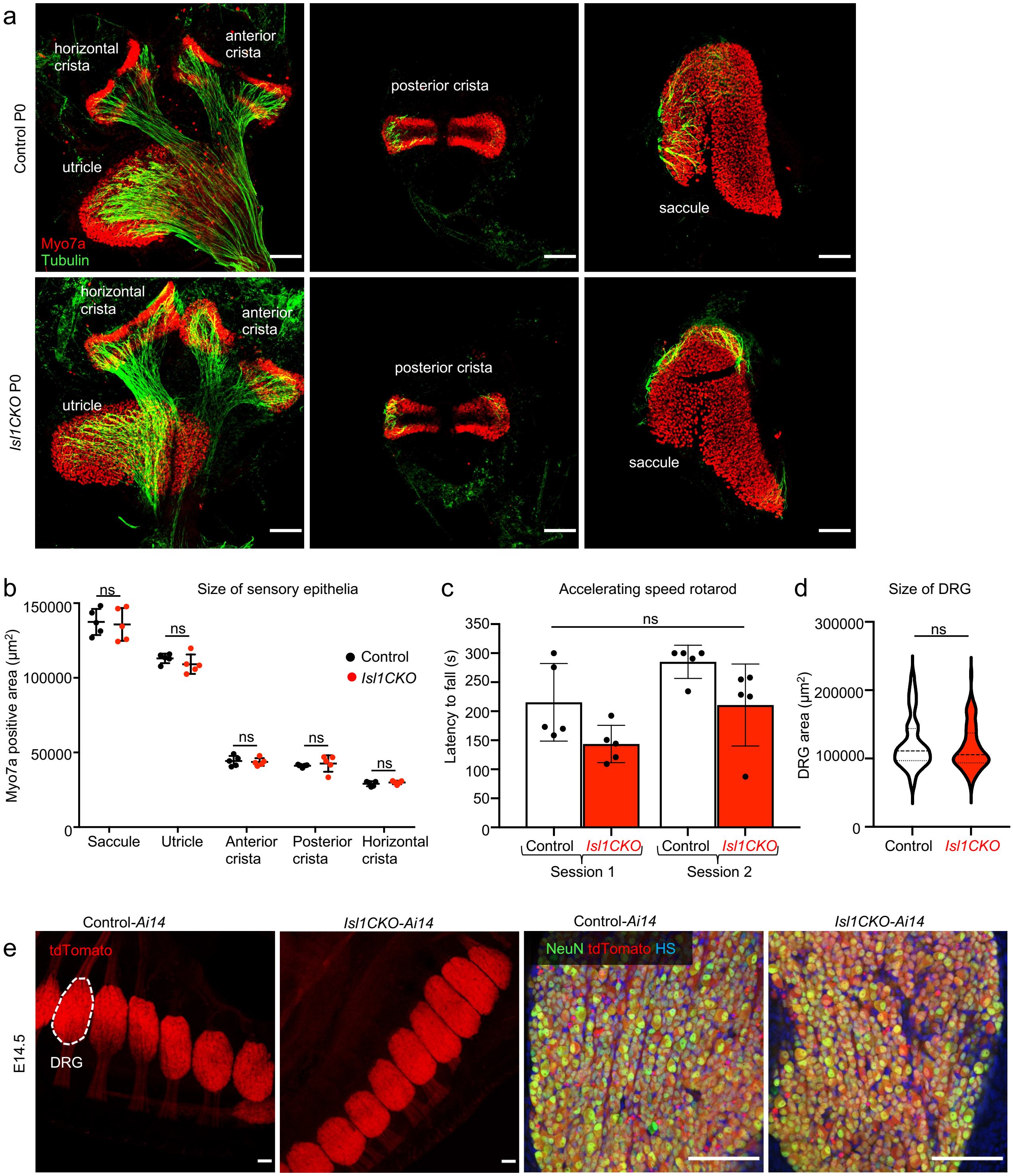


**Supplementary Fig. 8. Vestibular function unaffected in *Isl1CKO*. (a)** Representative images of whole-mount immunolabeling of the vestibular-end organs with anti-Myo7a (a marker of hair cells) and anti-α-tubulin (nerve fibers) at P0. **(b)** The relative size of Myo7a positive sensory epithelia of the vestibular-end organs is comparable between control and *Isl1CKO* mice. Error bars represent mean ± SD (n = 5 mice per genotype), *t*-tests (P > 0.05, ns, not significant). **(c)** Motor coordination of *Isl1CKO* is comparable to control mice, as evaluated on the accelerating rotarod. Mice were tested in three consecutive trials in one session per day. Histograms represent the means of three trials per animal and 5 mice per genotype ± SD, repeated measures ANOVA (P = 0.055, ns, not significant). **(d)** The quantification of the DRG area (n = 6 embryos per genotype and two sections per embryo). Violin plots indicate median (middle line), 25th, and 75th percentile (dotted lines). Mann-Whitney *U* test (P = 0.304, ns, not significant) **(e)** Representative images of sagittal sections of E14.5 embryos show normal-sized dorsal root ganglia (DRG) in *Isl1CKO*. Higher-magnification images show NeuN (a marker of differentiated neurons) and tdTomato positive DRG neurons. HS, Hoechst nuclear staining. Scale bars: 100 µm.

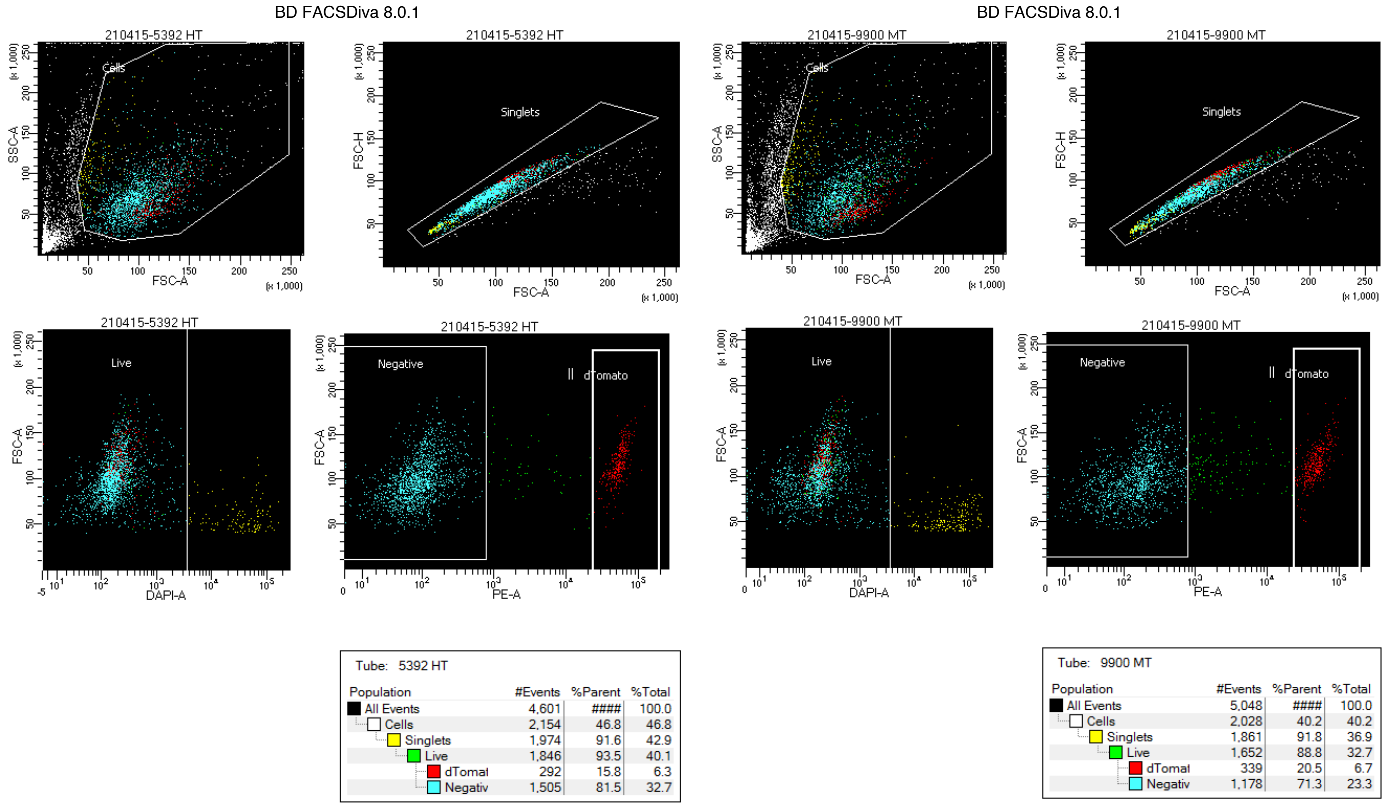


| Tube: 5392 HT | | | |
|---|---|---|---|
| Population | #Events | %Parent | %Total |
| All Events | 4,601 | #### | 100.0 |
| Cells | 2,154 | 46.8 | 46.8 |
| Singlets | 1,974 | 91.6 | 42.9 |
| Live | 1,846 | 93.5 | 40.1 |
| dTomat | 292 | 15.8 | 6.3 |
| Negativ | 1,505 | 81.5 | 32.7 |

| Tube: 9900 MT | | | |
|---|---|---|---|
| Population | #Events | %Parent | %Total |
| All Events | 5,048 | #### | 100.0 |
| Cells | 2,028 | 40.2 | 40.2 |
| Singlets | 1,861 | 91.8 | 36.9 |
| Live | 1,652 | 88.8 | 32.7 |
| dTomat | 339 | 20.5 | 6.7 |
| Negativ | 1,178 | 71.3 | 23.3 |

**Supplementary Fig. 9. Gating strategy used to isolate tdTomato$^+$ cells.** Representative example to show gating to purify live and individual tdTomato$^+$ cells for RNAseq.

**Supplementary Table 1.** Ribbon synapses (mean ± SD) per inner hair cell in the control and *Isl1CKO* cochlea of 2-month-old mice*

| Cochlear region | control | *Isl1CKO* |
|---|---|---|
| Base | 14.40 ± 3.346 | 6.57 ± 1.681 |
| Mid-base | 15.18 ± 3.154 | 7.54 ± 1.888 |
| Mid-apex | 13.65 ± 2.956 | 7.60 ± 2.618 |
| Apex | 11.04 ± 2.956 | 7.60 ± 2.068 |

*n = 6 cochlea per genotype/10 cells per each section (with the exception of only 5 samples of the apex in the control group)

### Supplementary Table 2. Primer sequences for RT-qPCR

| Gene name | Forward | Reverse | Product size |
|---|---|---|---|
| *Lhx1* | CCCATCCTGGACCGTTTCC | CGCTTGGAGAGATGCCCTG | 193 bp |
| *Lhx2* | CGAGGGCTCACGAAGACCAT | AGCAGGTAGTAGCGGTCAGA | 95 bp |
| *Cdh7* | AGCCAAAACGAGTTATACACTGC | TCTTAACCGTTGTTGTGTCACTG | 101 bp |
| *Nhlh2* | CAGCGTGTCGGACCTAGAG | CCAGGTTGAAAGCCTCCAC | 188 bp |
| *Ntrk2* | CTGGGGCTTATGCCTGCTG | AGGCTCAGTACACCAAATCCTA | 100 bp |
| *Ntrk3* | CTGAGTGCTACAATCTAAGCCC | CACACCCCATAGAACTTGACAAT | 157 bp |
| *Dcc* | GCTATGGTGTTGGCAGTCCT | TAATCAACGGGGTCAGTGGG | 196 bp |
| *Epha5* | CTGGCGGACGGAAAGATGT | TTCAGGCCGATTTGCTGGG | 118 bp |
| *Gata3* | CCTCTACGCTCCTTGCTACTC | AGAGGAATCCGAGTGTGACC | 94 bp |
| *Ntng1* | CCAGTATCGGTACTAATGTCTGC | CCGTAGCTTCTCACAGAGGATG | 136 bp |
| *Prdm8* | AGAACGCCATATTCGGTCCC | ATTTGCCGCCGAAGTGTCTA | 131 bp |
| *Robo2* | GCTGAGAATCGGGTGGGAAA | AACTGTGGAGGAGCAACAGG | 77 bp |
| *Slit2* | CGAGAGTTTGTCTGCAGTGATG | CTACAGGTACAAGCAGCGGG | 95 bp |
| *Tbx3* | CAACTCTCGGTGGATGGTGG | TTGCGTGATCGCTTGGGAA | 188 bp |
| *Uncx* | ACCCGCACCAACTTTACCG | TGAACTCGGGACTCGACCA | 128 bp |
| *Hprt* | GCTTGCTGGTGAAAAGGACCTCTCGAAG | CCTGAAGTACTCATTATAGTCAAGGGCAT | 117 bp |
